# SKIN REINNERVATION BY COLLATERAL SPROUTING FOLLOWING SPARED NERVE INJURY IN MICE

**DOI:** 10.1101/2023.09.12.557420

**Authors:** S.M. Jeon, A. Pradeep, D. Chang, L. McDonough, Y. Chen, A. Latremoliere, L.K. Crawford, M.J. Caterina

## Abstract

Following peripheral nerve injury, denervated tissues can be reinnervated via regeneration of injured neurons or via collateral sprouting of neighboring uninjured afferents into the denervated territory. While there has been substantial focus on mechanisms underlying regeneration, collateral sprouting has received relatively less attention. In this study, we used immunohistochemistry and genetic neuronal labeling to define the subtype specificity of sprouting-mediated reinnervation of plantar hind paw skin in the mouse spared nerve injury (SNI) model, in which productive regeneration cannot occur. Following an initial loss of cutaneous afferents in the tibial nerve territory, we observed progressive centripetal reinnervation by multiple subtypes of neighboring uninjured fibers into denervated glabrous and hairy plantar skin. In addition to dermal reinnervation, CGRP-expressing peptidergic fibers slowly but continuously repopulated the denervated epidermis, Interestingly, GFRα2-expressing nonpeptidergic fibers exhibited a transient burst of epidermal reinnervation, followed by trend towards regression. Presumptive sympathetic nerve fibers also sprouted into the denervated territory, as did a population of myelinated TrkC lineage fibers, though the latter did so less efficiently. Conversely, rapidly adapting Aβ fiber and C fiber low threshold mechanoreceptor (LTMR) subtypes failed to exhibit convincing collateral sprouting up to 8 weeks after nerve injury. Optogenetics and behavioral assays further demonstrated the functionality of collaterally sprouted fibers in hairy plantar skin with restoration of punctate mechanosensation without hypersensitivity. Our findings advance understanding of differential collateral sprouting among sensory neuron subpopulations and may guide strategies to promote the progression of sensory recovery or limit maladaptive sensory phenomena after peripheral nerve injury.

**Significance Statement:** Following nerve injury, whereas one mechanism for tissue reinnervation is regeneration of injured neurons, another, less well studied mechanism is collateral sprouting of nearby uninjured neurons. In this study, we examined collateral sprouting in denervated mouse skin and showed that it involves some, but not all neuronal subtypes. Despite such heterogeneity, a significant degree of restoration of punctate mechanical sensitivity is achieved. These findings highlight the diversity of collateral sprouting among peripheral neuron subtypes and reveal important differences between pre- and post-denervation skin that might be appealing targets for therapeutic correction to enhance functional recovery from denervation and prevent unwanted sensory phenomena such as pain or numbness.

## Introduction

Injury to peripheral nerves can not only impair motor function but also interfere with sensory functions necessary for protective reflexes and tactile acuity. Furthermore, nerve injury can lead to abnormal neuronal excitability and consequent neuropathic pain (Ji and Strichartz, 2004; Campbell and Meyer, 2006; Costigan et al., 2009). Recovery from nerve injury occurs via two distinct processes. In the first of these, regeneration, injured axons grow along their original paths to reinnervate target tissues (Rigoni and Negro, 2020). While this represents one potential means of achieving restoration of sensory function, it is sometimes impossible, due to an excessive gap in the injured nerve. Even when regeneration is possible, if a nerve is injured far from its target tissues, the inherently slow rate of regeneration delays and might even preclude functional restoration (Nath and Mackinnon 2000). A second mechanism of reinnervation, collateral sprouting, involves local ectopic branching and axonal extension from uninjured (i.e., spared) neurons into adjacent denervated tissue (Livingston, 1947; Ahcan et al., 1998; Theriault et al., 1998; Ali et al., 2002). Because it is initiated locally, collateral sprouting theoretically offers the opportunity for a more rapid sensory recovery than nerve regeneration, and could compensate when regeneration is impossible or delayed (Ahcan et al., 1998). However, available data suggest that collateral sprouting in humans is limited in extent, and that in both humans and in animal models, collateral sprouting is nonuniform across sensory modalities and neuronal subtypes (Horch, 1981; Jackson and Diamond, 1984a; Ahcan et al., 1998; Diamond and Foerster, 1992). There is also a complex relationship between tissue reinnervation and the duration of neuropathic pain, and in some cases neuropathic pain may occur in regions of collateral sprouting (Ali et al., 2002; Gangadharan et al., 2022). Given these considerations, elucidation of mechanisms governing both regeneration and collateral sprouting is necessary to facilitate the restoration of desirable sensory function and may help avoid or reverse the development of post-injury pathological pain.

In this study, we used whole-mount tissue staining, genetically labeled mouse lines, and both conventional and optogenetic behavioral assays to anatomically and functionally assess collateral sprouting in the mouse spared nerve injury model. Following initial denervation, we observed significant centripetal sprouting by some, but not all subpopulations of sensory neurons assayed. Moreover, while peptidergic nociceptors exhibited a slow, steady timecourse of epidermal reinnervation, nonpeptidergic fibers showed a transient burst of such reinnervation that subsequently regressed. Meanwhile, we observed a paucity or lack of sprouting among some low-threshold mechanoreceptor subtypes. Despite such heterogeneity, we observed apparently faithful restoration of punctate mechanosensitivity across a wide range of forces in this skin territory without evident overshoot. Together, our data highlight the subtype specificity of collateral sprouting and illustrate the sufficiency of those fibers that do sprout to restore significant sensitivity to denervated hind paw skin.

## Materials and Methods

### Mouse Strains

TH^2ACreER^, Split^Cre^, TrkC^tdTomato^, TrkC^CreER^ mouse lines(Li et al., 2011; Abraira et al., 2017; Bai et al., 2015) were generously provided by Dr. David Ginty. Pirt^Cre^ mice(Kim et al., 2014) were kindly provided by Dr. Xinzhong Dong. Rosa26^LSLEYFP^ (Ai3, #007903), Rosa26^LSLtdTomato^ (Ai9, #007909), Rosa26^LSL-ChR2-EYFP^ (Ai32, #024109) and C57BL/6J (#000664) mice were obtained from the Jackson Laboratory. Separate groups of mice at least 7-8 weeks old at the start of the experiments, were used for behavioral and anatomical studies, respectively. Age-matched mice from the same or parallel mating cages were randomly assigned to experimental groups in injured vs. sham comparisons. Mice were housed 1 to 5 per cage on a 14hr light/10hr dark light cycle, and provided food and water ad libitum. Mice were handled in accordance with the Johns Hopkins University Institutional Animal Care and Use Committee guidelines as well as National Institutes of Health Guide for the Care and Use of Laboratory Animals.

### Peripheral nerve Injury model

Spared nerve injury (SNI) was performed as previously described (Bourquin et al., 2006). In brief, under deep isoflurane anesthesia, one sciatic nerve of mice 7-8 weeks old was exposed in the thigh region, near its trifurcation, and the tibial and common peroneal nerve branches were ligated. A small section immediately distal to the ligation was excised. The sural nerve branch was left intact, avoiding contact or stretching. Finally, muscle and skin were sutured in two distinct layers with silk 6-0 and 4-0 sutures, respectively. In a limited number of mice, a second surgical exposure of the sciatic nerve was performed 8 weeks after sural sparing SNI and the remaining sural branch was transected, a small piece distal to the transection was excised, and muscle and skin were again closed in two layers.

### Behavioral Testing

Experiments were performed at baseline (usually the day before surgery) and up to 2 months after SNI surgery. For assays using pinprick, 1.4 g von Frey filament, and optogenetic stimulation, experiments were performed at baseline and at intervals of 3 or 4 days for one month after SNI. For assays with von Frey filaments over the full range of forces, experiments were performed at baseline and at 1, 4, and 8 weeks after SNI. For assays using the 0.4 g von Frey filament, experiments were performed at baseline and weekly for 8wks after SNI. Behavioral assays for optogenetics were conducted with the experimenter blinded to genotype or to sham vs. SNI treatment. Animal numbers for each experiment are indicated in the figures and in Table 1.

**Table 1.**
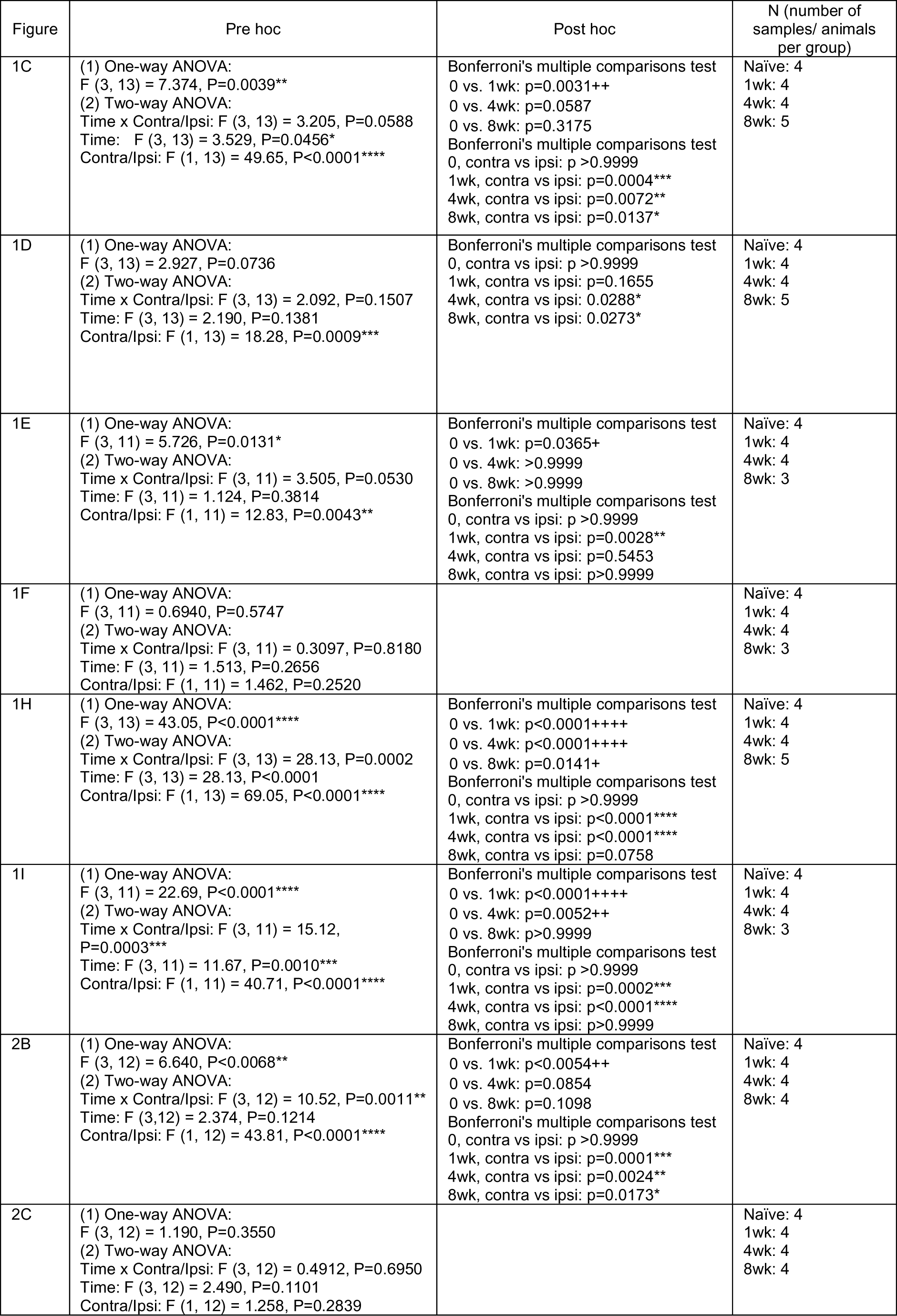
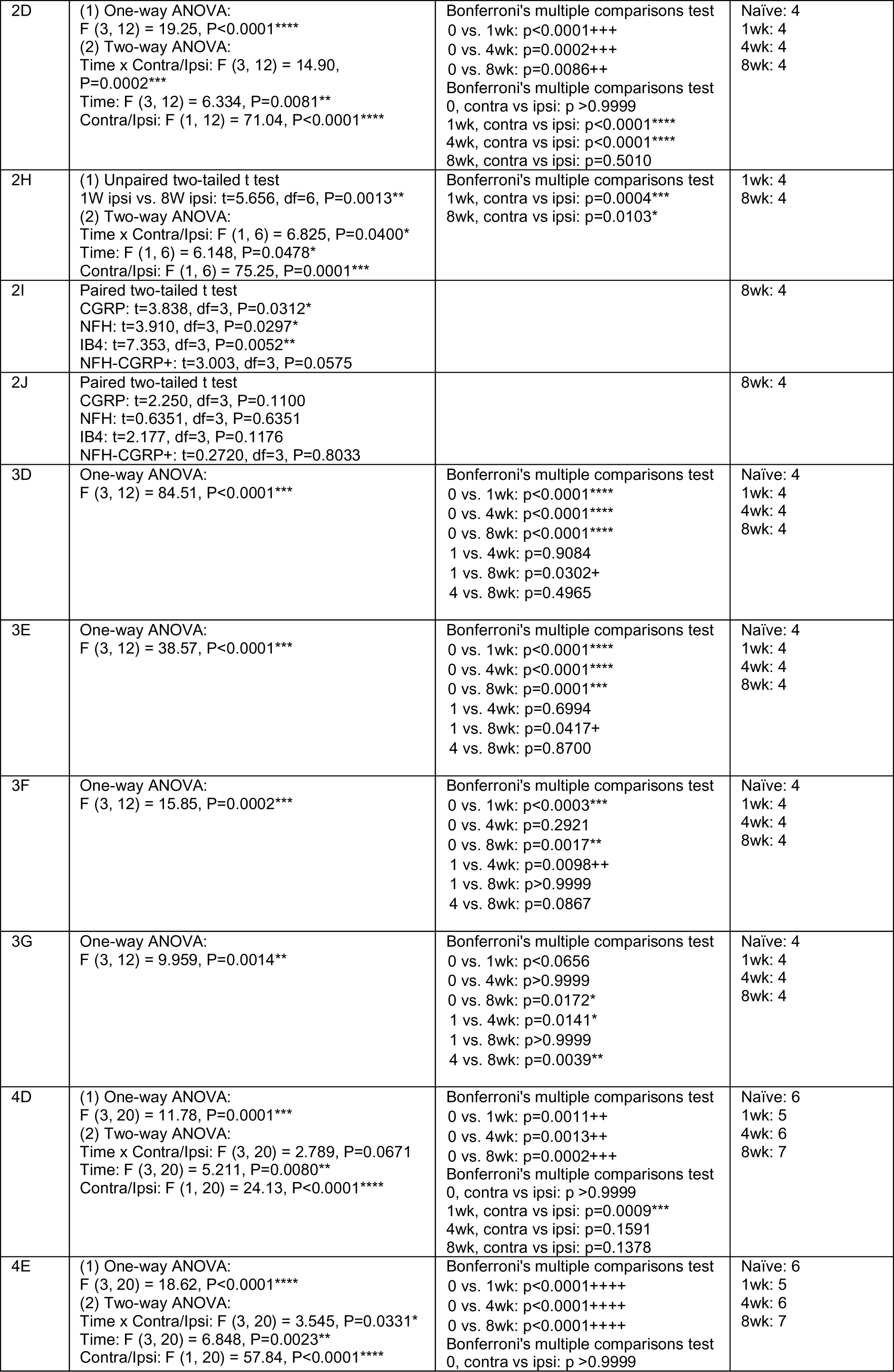
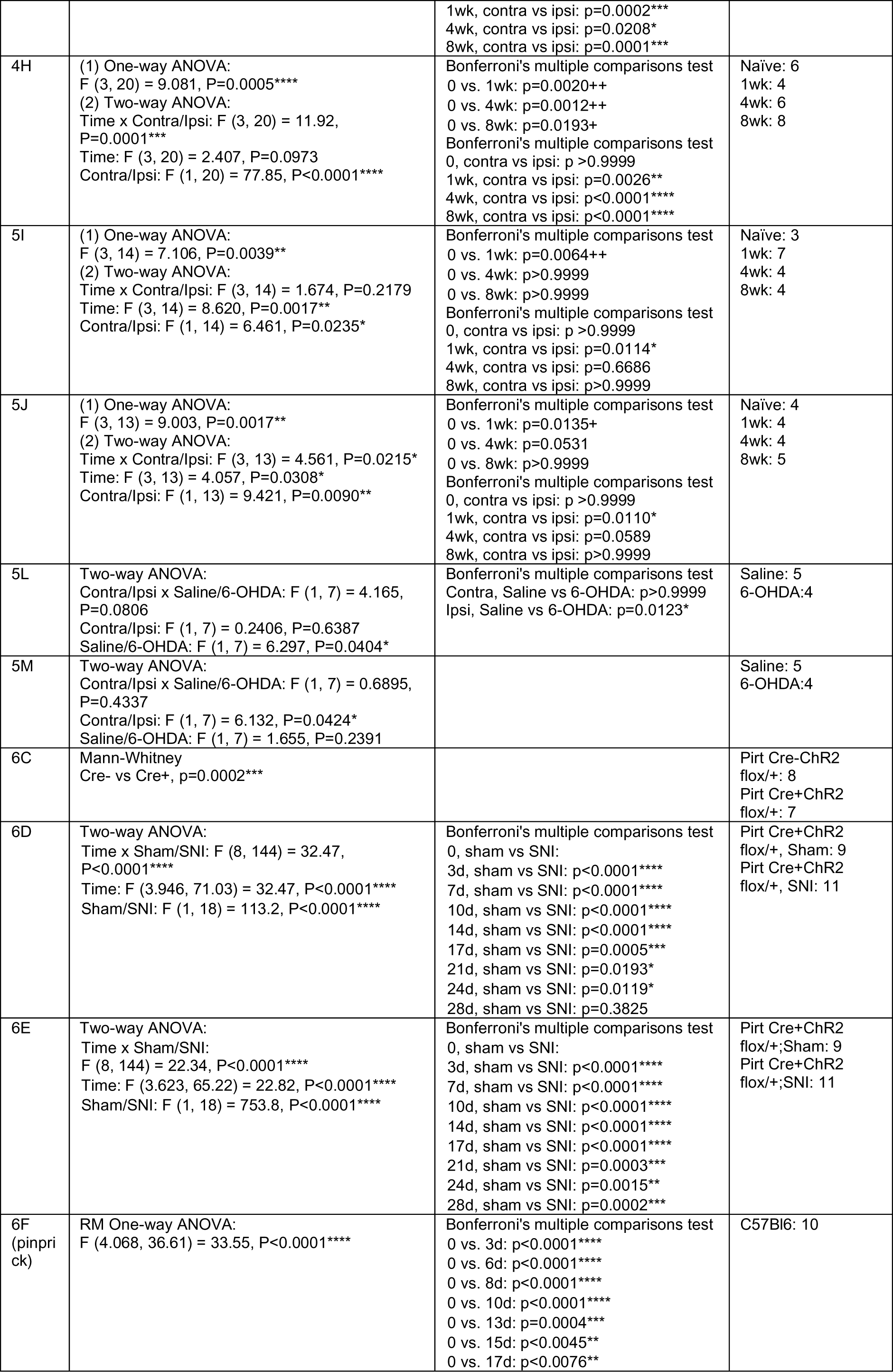
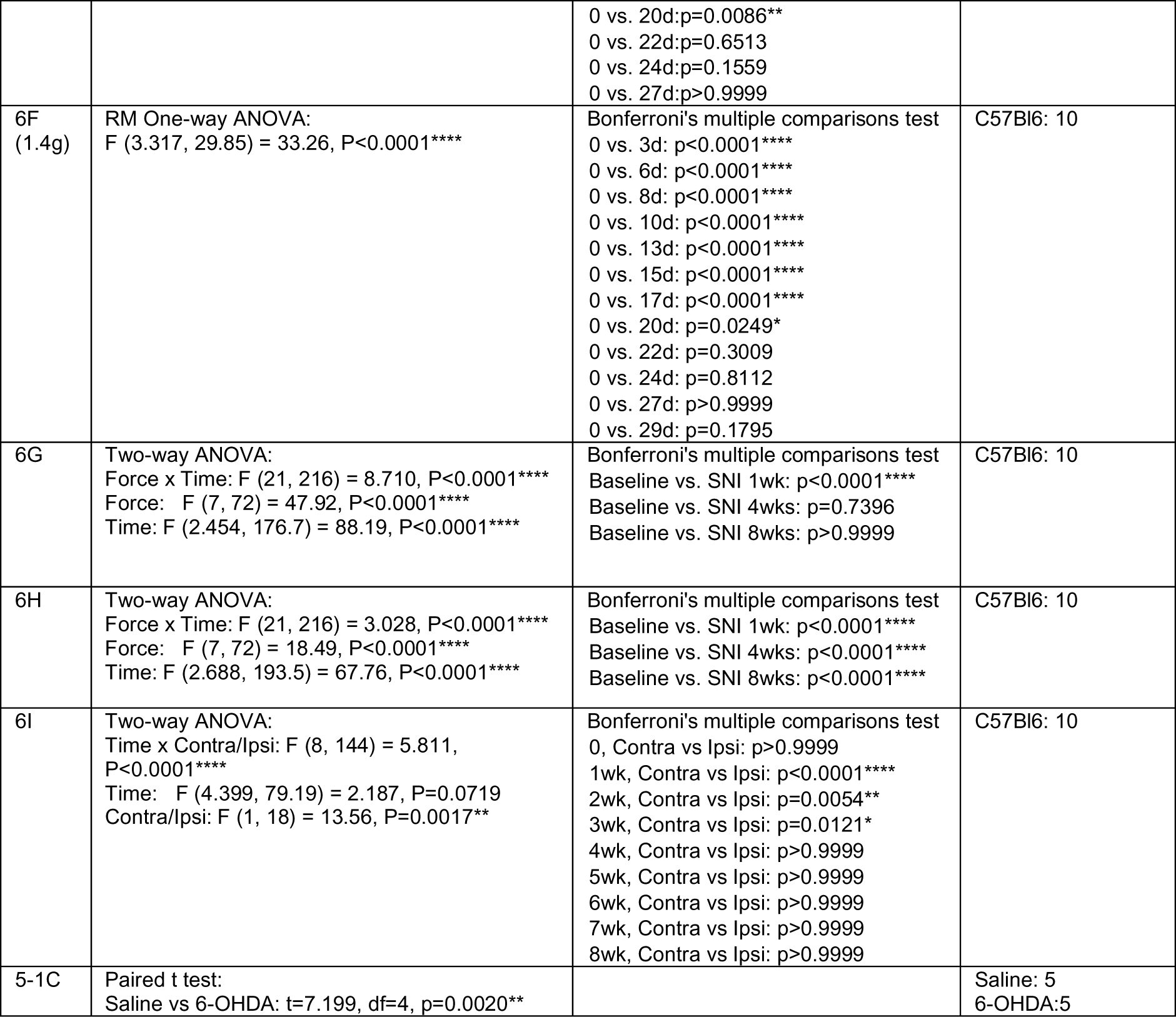
Statistical analysis and the number of animals/samples used in the experiments.

| Figure | Pre hoc | Post hoc | N (number of samples/ animals per group) |
| --- | --- | --- | --- |
| 1C | (1) One-way ANOVA:<br>F (3, 13) = 7.374, P=0.0039**<br>(2) Two-way ANOVA:<br>Time x Contra/Ipsi: F (3, 13) = 3.205, P=0.0588<br>Time: F (3, 13) = 3.529, P=0.0456*<br>Contra/Ipsi: F (1, 13) = 49.65, P<0.0001**** | Bonferroni's multiple comparisons test<br>0 vs. 1wk: p=0.0031++<br>0 vs. 4wk: p=0.0587<br>0 vs. 8wk: p=0.3175<br>Bonferroni's multiple comparisons test<br>0, contra vs ipsi: p >0.9999<br>1wk, contra vs ipsi: p=0.0004***<br>4wk, contra vs ipsi: p=0.0072**<br>8wk, contra vs ipsi: p=0.0137* | Naïve: 4<br>1wk: 4<br>4wk: 4<br>8wk: 5 |
| 1D | (1) One-way ANOVA:<br>F (3, 13) = 2.927, P=0.0736<br>(2) Two-way ANOVA:<br>Time x Contra/Ipsi: F (3, 13) = 2.092, P=0.1507<br>Time: F (3, 13) = 2.190, P=0.1381<br>Contra/Ipsi: F (1, 13) = 18.28, P=0.0009*** | Bonferroni's multiple comparisons test<br>0, contra vs ipsi: p >0.9999<br>1wk, contra vs ipsi: p=0.1655<br>4wk, contra vs ipsi: 0.0288*<br>8wk, contra vs ipsi: 0.0273* | Naïve: 4<br>1wk: 4<br>4wk: 4<br>8wk: 5 |
| 1E | (1) One-way ANOVA:<br>F (3, 11) = 5.726, P=0.0131*<br>(2) Two-way ANOVA:<br>Time x Contra/Ipsi: F (3, 11) = 3.505, P=0.0530<br>Time: F (3, 11) = 1.124, P=0.3814<br>Contra/Ipsi: F (1, 11) = 12.83, P=0.0043** | Bonferroni's multiple comparisons test<br>0 vs. 1wk: p=0.0365+<br>0 vs. 4wk: >0.9999<br>0 vs. 8wk: >0.9999<br>Bonferroni's multiple comparisons test<br>0, contra vs ipsi: p >0.9999<br>1wk, contra vs ipsi: p=0.0028**<br>4wk, contra vs ipsi: p=0.5453<br>8wk, contra vs ipsi: p>0.9999 | Naïve: 4<br>1wk: 4<br>4wk: 4<br>8wk: 3 |
| 1F | (1) One-way ANOVA:<br>F (3, 11) = 0.6940, P=0.5747<br>(2) Two-way ANOVA:<br>Time x Contra/Ipsi: F (3, 11) = 0.3097, P=0.8180<br>Time: F (3, 11) = 1.513, P=0.2656<br>Contra/Ipsi: F (1, 11) = 1.462, P=0.2520 |  | Naïve: 4<br>1wk: 4<br>4wk: 4<br>8wk: 3 |
| 1H | (1) One-way ANOVA:<br>F (3, 13) = 43.05, P<0.0001****<br>(2) Two-way ANOVA:<br>Time x Contra/Ipsi: F (3, 13) = 28.13, P=0.0002<br>Time: F (3, 13) = 28.13, P<0.0001<br>Contra/Ipsi: F (1, 13) = 69.05, P<0.0001**** | Bonferroni's multiple comparisons test<br>0 vs. 1wk: p<0.0001++++<br>0 vs. 4wk: p<0.0001++++<br>0 vs. 8wk: p=0.0141+<br>Bonferroni's multiple comparisons test<br>0, contra vs ipsi: p >0.9999<br>1wk, contra vs ipsi: p<0.0001****<br>4wk, contra vs ipsi: p<0.0001****<br>8wk, contra vs ipsi: p=0.0758 | Naïve: 4<br>1wk: 4<br>4wk: 4<br>8wk: 5 |
| 1I | (1) One-way ANOVA:<br>F (3, 11) = 22.69, P<0.0001****<br>(2) Two-way ANOVA:<br>Time x Contra/Ipsi: F (3, 11) = 15.12, P=0.0003***<br>Time: F (3, 11) = 11.67, P=0.0010***<br>Contra/Ipsi: F (1, 11) = 40.71, P<0.0001**** | Bonferroni's multiple comparisons test<br>0 vs. 1wk: p<0.0001++++<br>0 vs. 4wk: p=0.0052++<br>0 vs. 8wk: p>0.9999<br>Bonferroni's multiple comparisons test<br>0, contra vs ipsi: p >0.9999<br>1wk, contra vs ipsi: p=0.0002***<br>4wk, contra vs ipsi: p<0.0001****<br>8wk, contra vs ipsi: p>0.9999 | Naïve: 4<br>1wk: 4<br>4wk: 4<br>8wk: 3 |
| 2B | (1) One-way ANOVA:<br>F (3, 12) = 6.640, P<0.0068**<br>(2) Two-way ANOVA:<br>Time x Contra/Ipsi: F (3, 12) = 10.52, P=0.0011**<br>Time: F (3, 12) = 2.374, P=0.1214<br>Contra/Ipsi: F (1, 12) = 43.81, P<0.0001**** | Bonferroni's multiple comparisons test<br>0 vs. 1wk: p<0.0054++<br>0 vs. 4wk: p=0.0854<br>0 vs. 8wk: p=0.1098<br>Bonferroni's multiple comparisons test<br>0, contra vs ipsi: p >0.9999<br>1wk, contra vs ipsi: p=0.0001***<br>4wk, contra vs ipsi: p=0.0024**<br>8wk, contra vs ipsi: p=0.0173* | Naïve: 4<br>1wk: 4<br>4wk: 4<br>8wk: 4 |
| 2C | (1) One-way ANOVA:<br>F (3, 12) = 1.190, P=0.3550<br>(2) Two-way ANOVA:<br>Time x Contra/Ipsi: F (3, 12) = 0.4912, P=0.6950<br>Time: F (3, 12) = 2.490, P=0.1101<br>Contra/Ipsi: F (1, 12) = 1.258, P=0.2839 |  | Naïve: 4<br>1wk: 4<br>4wk: 4<br>8wk: 4 |
| 2D | (1) One-way ANOVA:<br>F (3, 12) = 19.25, P<0.0001****<br>(2) Two-way ANOVA:<br>Time x Contra/Ipsi: F (3, 12) = 14.90,<br>P=0.0002***<br>Time: F (3, 12) = 6.334, P=0.0081**<br>Contra/Ipsi: F (1, 12) = 71.04, P<0.0001**** | Bonferroni's multiple comparisons test<br>0 vs. 1wk: p<0.0001+++<br>0 vs. 4wk: p=0.0002+++<br>0 vs. 8wk: p=0.0086++<br>Bonferroni's multiple comparisons test<br>0, contra vs ipsi: p >0.9999<br>1wk, contra vs ipsi: p<0.0001****<br>4wk, contra vs ipsi: p<0.0001****<br>8wk, contra vs ipsi: p=0.5010 | Naïve: 4<br>1wk: 4<br>4wk: 4<br>8wk: 4 |
| 2H | (1) Unpaired two-tailed t test<br>1W ipsi vs. 8W ipsi: t=5.656, df=6, P=0.0013**<br>(2) Two-way ANOVA:<br>Time x Contra/Ipsi: F (1, 6) = 6.825, P=0.0400*<br>Time: F (1, 6) = 6.148, P=0.0478*<br>Contra/Ipsi: F (1, 6) = 75.25, P=0.0001*** | Bonferroni's multiple comparisons test<br>1wk, contra vs ipsi: p=0.0004***<br>8wk, contra vs ipsi: p=0.0103* | 1wk: 4<br>8wk: 4 |
| 2I | Paired two-tailed t test<br>CGRP: t=3.838, df=3, P=0.0312*<br>NFH: t=3.910, df=3, P=0.0297*<br>IB4: t=7.353, df=3, P=0.0052**<br>NFH-CGRP+: t=3.003, df=3, P=0.0575 |  | 8wk: 4 |
| 2J | Paired two-tailed t test<br>CGRP: t=2.250, df=3, P=0.1100<br>NFH: t=0.6351, df=3, P=0.6351<br>IB4: t=2.177, df=3, P=0.1176<br>NFH-CGRP+: t=0.2720, df=3, P=0.8033 |  | 8wk: 4 |
| 3D | One-way ANOVA:<br>F (3, 12) = 84.51, P<0.0001*** | Bonferroni's multiple comparisons test<br>0 vs. 1wk: p<0.0001****<br>0 vs. 4wk: p<0.0001****<br>0 vs. 8wk: p<0.0001****<br>1 vs. 4wk: p=0.9084<br>1 vs. 8wk: p=0.0302+<br>4 vs. 8wk: p=0.4965 | Naïve: 4<br>1wk: 4<br>4wk: 4<br>8wk: 4 |
| 3E | One-way ANOVA:<br>F (3, 12) = 38.57, P<0.0001*** | Bonferroni's multiple comparisons test<br>0 vs. 1wk: p<0.0001****<br>0 vs. 4wk: p<0.0001****<br>0 vs. 8wk: p=0.0001***<br>1 vs. 4wk: p=0.6994<br>1 vs. 8wk: p=0.0417+<br>4 vs. 8wk: p=0.8700 | Naïve: 4<br>1wk: 4<br>4wk: 4<br>8wk: 4 |
| 3F | One-way ANOVA:<br>F (3, 12) = 15.85, P=0.0002*** | Bonferroni's multiple comparisons test<br>0 vs. 1wk: p<0.0003***<br>0 vs. 4wk: p=0.2921<br>0 vs. 8wk: p=0.0017**<br>1 vs. 4wk: p=0.0098++<br>1 vs. 8wk: p>0.9999<br>4 vs. 8wk: p=0.0867 | Naïve: 4<br>1wk: 4<br>4wk: 4<br>8wk: 4 |
| 3G | One-way ANOVA:<br>F (3, 12) = 9.959, P=0.0014** | Bonferroni's multiple comparisons test<br>0 vs. 1wk: p<0.0656<br>0 vs. 4wk: p>0.9999<br>0 vs. 8wk: p=0.0172*<br>1 vs. 4wk: p=0.0141*<br>1 vs. 8wk: p>0.9999<br>4 vs. 8wk: p=0.0039** | Naïve: 4<br>1wk: 4<br>4wk: 4<br>8wk: 4 |
| 4D | (1) One-way ANOVA:<br>F (3, 20) = 11.78, P=0.0001***<br>(2) Two-way ANOVA:<br>Time x Contra/Ipsi: F (3, 20) = 2.789, P=0.0671<br>Time: F (3, 20) = 5.211, P=0.0080**<br>Contra/Ipsi: F (1, 20) = 24.13, P<0.0001**** | Bonferroni's multiple comparisons test<br>0 vs. 1wk: p=0.0011++<br>0 vs. 4wk: p=0.0013++<br>0 vs. 8wk: p=0.0002+++<br>Bonferroni's multiple comparisons test<br>0, contra vs ipsi: p >0.9999<br>1wk, contra vs ipsi: p=0.0009***<br>4wk, contra vs ipsi: p=0.1591<br>8wk, contra vs ipsi: p=0.1378 | Naïve: 6<br>1wk: 5<br>4wk: 6<br>8wk: 7 |
| 4E | (1) One-way ANOVA:<br>F (3, 20) = 18.62, P<0.0001****<br>(2) Two-way ANOVA:<br>Time x Contra/Ipsi: F (3, 20) = 3.545, P=0.0331*<br>Time: F (3, 20) = 6.848, P=0.0023**<br>Contra/Ipsi: F (1, 20) = 57.84, P<0.0001**** | Bonferroni's multiple comparisons test<br>0 vs. 1wk: p<0.0001++++<br>0 vs. 4wk: p<0.0001++++<br>0 vs. 8wk: p<0.0001++++<br>Bonferroni's multiple comparisons test<br>0, contra vs ipsi: p >0.9999 | Naïve: 6<br>1wk: 5<br>4wk: 6<br>8wk: 7 |
|  |  | 1wk, contra vs ipsi: p=0.0002***<br>4wk, contra vs ipsi: p=0.0208*<br>8wk, contra vs ipsi: p=0.0001*** |  |
| 4H | (1) One-way ANOVA:<br>F (3, 20) = 9.081, P=0.0005****<br>(2) Two-way ANOVA:<br>Time x Contra/Ipsi: F (3, 20) = 11.92, P=0.0001***<br>Time: F (3, 20) = 2.407, P=0.0973<br>Contra/Ipsi: F (1, 20) = 77.85, P<0.0001**** | Bonferroni's multiple comparisons test<br>0 vs. 1wk: p=0.0020++<br>0 vs. 4wk: p=0.0012++<br>0 vs. 8wk: p=0.0193+<br>Bonferroni's multiple comparisons test<br>0, contra vs ipsi: p>0.9999<br>1wk, contra vs ipsi: p=0.0026**<br>4wk, contra vs ipsi: p<0.0001****<br>8wk, contra vs ipsi: p<0.0001**** | Naïve: 6<br>1wk: 4<br>4wk: 6<br>8wk: 8 |
| 5I | (1) One-way ANOVA:<br>F (3, 14) = 7.106, P=0.0039**<br>(2) Two-way ANOVA:<br>Time x Contra/Ipsi: F (3, 14) = 1.674, P=0.2179<br>Time: F (3, 14) = 8.620, P=0.0017**<br>Contra/Ipsi: F (1, 14) = 6.461, P=0.0235* | Bonferroni's multiple comparisons test<br>0 vs. 1wk: p=0.0064++<br>0 vs. 4wk: p>0.9999<br>0 vs. 8wk: p>0.9999<br>Bonferroni's multiple comparisons test<br>0, contra vs ipsi: p>0.9999<br>1wk, contra vs ipsi: p=0.0114*<br>4wk, contra vs ipsi: p=0.6686<br>8wk, contra vs ipsi: p>0.9999 | Naïve: 3<br>1wk: 7<br>4wk: 4<br>8wk: 4 |
| 5J | (1) One-way ANOVA:<br>F (3, 13) = 9.003, P=0.0017**<br>(2) Two-way ANOVA:<br>Time x Contra/Ipsi: F (3, 13) = 4.561, P=0.0215*<br>Time: F (3, 13) = 4.057, P=0.0308*<br>Contra/Ipsi: F (1, 13) = 9.421, P=0.0090** | Bonferroni's multiple comparisons test<br>0 vs. 1wk: p=0.0135+<br>0 vs. 4wk: p=0.0531<br>0 vs. 8wk: p>0.9999<br>Bonferroni's multiple comparisons test<br>0, contra vs ipsi: p>0.9999<br>1wk, contra vs ipsi: p=0.0110*<br>4wk, contra vs ipsi: p=0.0589<br>8wk, contra vs ipsi: p>0.9999 | Naïve: 4<br>1wk: 4<br>4wk: 4<br>8wk: 5 |
| 5L | Two-way ANOVA:<br>Contra/Ipsi x Saline/6-OHDA: F (1, 7) = 4.165, P=0.0806<br>Contra/Ipsi: F (1, 7) = 0.2406, P=0.6387<br>Saline/6-OHDA: F (1, 7) = 6.297, P=0.0404* | Bonferroni's multiple comparisons test<br>Contra, Saline vs 6-OHDA: p>0.9999<br>Ipsi, Saline vs 6-OHDA: p=0.0123* | Saline: 5<br>6-OHDA: 4 |
| 5M | Two-way ANOVA:<br>Contra/Ipsi x Saline/6-OHDA: F (1, 7) = 0.6895, P=0.4337<br>Contra/Ipsi: F (1, 7) = 6.132, P=0.0424*<br>Saline/6-OHDA: F (1, 7) = 1.655, P=0.2391 |  | Saline: 5<br>6-OHDA: 4 |
| 6C | Mann-Whitney<br>Cre- vs Cre+, p=0.0002*** |  | Pirt Cre-ChR2<br>flox/+: 8<br>Pirt Cre+ChR2<br>flox/+: 7 |
| 6D | Two-way ANOVA:<br>Time x Sham/SNI: F (8, 144) = 32.47, P<0.0001****<br>Time: F (3.946, 71.03) = 32.47, P<0.0001****<br>Sham/SNI: F (1, 18) = 113.2, P<0.0001**** | Bonferroni's multiple comparisons test<br>0, sham vs SNI:<br>3d, sham vs SNI: p<0.0001****<br>7d, sham vs SNI: p<0.0001****<br>10d, sham vs SNI: p<0.0001****<br>14d, sham vs SNI: p<0.0001****<br>17d, sham vs SNI: p=0.0005***<br>21d, sham vs SNI: p=0.0193*<br>24d, sham vs SNI: p=0.0119*<br>28d, sham vs SNI: p=0.3825 | Pirt Cre+ChR2<br>flox/+, Sham: 9<br>Pirt Cre+ChR2<br>flox/+, SNI: 11 |
| 6E | Two-way ANOVA:<br>Time x Sham/SNI:<br>F (8, 144) = 22.34, P<0.0001****<br>Time: F (3.623, 65.22) = 22.82, P<0.0001****<br>Sham/SNI: F (1, 18) = 753.8, P<0.0001**** | Bonferroni's multiple comparisons test<br>0, sham vs SNI:<br>3d, sham vs SNI: p<0.0001****<br>7d, sham vs SNI: p<0.0001****<br>10d, sham vs SNI: p<0.0001****<br>14d, sham vs SNI: p<0.0001****<br>17d, sham vs SNI: p<0.0001****<br>21d, sham vs SNI: p=0.0003***<br>24d, sham vs SNI: p=0.0015**<br>28d, sham vs SNI: p=0.0002*** | Pirt Cre+ChR2<br>flox/+, Sham: 9<br>Pirt Cre+ChR2<br>flox/+, SNI: 11 |
| 6F<br>(pinprick) | RM One-way ANOVA:<br>F (4.068, 36.61) = 33.55, P<0.0001**** | Bonferroni's multiple comparisons test<br>0 vs. 3d: p<0.0001****<br>0 vs. 6d: p<0.0001****<br>0 vs. 8d: p<0.0001****<br>0 vs. 10d: p<0.0001****<br>0 vs. 13d: p=0.0004***<br>0 vs. 15d: p<0.0045**<br>0 vs. 17d: p<0.0076** | C57Bl6: 10 |
| | | 0 vs. 20d: $p=0.0086^{**}$<br>0 vs. 22d: $p=0.6513$<br>0 vs. 24d: $p=0.1559$<br>0 vs. 27d: $p>0.9999$ | |
| 6F<br>(1.4g) | RM One-way ANOVA:<br>$F(3.317, 29.85) = 33.26, P<0.0001^{****}$ | Bonferroni's multiple comparisons test<br>0 vs. 3d: $p<0.0001^{****}$<br>0 vs. 6d: $p<0.0001^{****}$<br>0 vs. 8d: $p<0.0001^{****}$<br>0 vs. 10d: $p<0.0001^{****}$<br>0 vs. 13d: $p<0.0001^{****}$<br>0 vs. 15d: $p<0.0001^{****}$<br>0 vs. 17d: $p<0.0001^{****}$<br>0 vs. 20d: $p=0.0249^{*}$<br>0 vs. 22d: $p=0.3009$<br>0 vs. 24d: $p=0.8112$<br>0 vs. 27d: $p>0.9999$<br>0 vs. 29d: $p=0.1795$ | C57Bl6: 10 |
| 6G | Two-way ANOVA:<br>Force x Time: $F(21, 216) = 8.710, P<0.0001^{****}$<br>Force: $F(7, 72) = 47.92, P<0.0001^{****}$<br>Time: $F(2.454, 176.7) = 88.19, P<0.0001^{****}$ | Bonferroni's multiple comparisons test<br>Baseline vs. SNI 1wk: $p<0.0001^{****}$<br>Baseline vs. SNI 4wks: $p=0.7396$<br>Baseline vs. SNI 8wks: $p>0.9999$ | C57Bl6: 10 |
| 6H | Two-way ANOVA:<br>Force x Time: $F(21, 216) = 3.028, P<0.0001^{****}$<br>Force: $F(7, 72) = 18.49, P<0.0001^{****}$<br>Time: $F(2.688, 193.5) = 67.76, P<0.0001^{****}$ | Bonferroni's multiple comparisons test<br>Baseline vs. SNI 1wk: $p<0.0001^{****}$<br>Baseline vs. SNI 4wks: $p<0.0001^{****}$<br>Baseline vs. SNI 8wks: $p<0.0001^{****}$ | C57Bl6: 10 |
| 6I | Two-way ANOVA:<br>Time x Contra/Ipsi: $F(8, 144) = 5.811, P<0.0001^{****}$<br>Time: $F(4.399, 79.19) = 2.187, P=0.0719$<br>Contra/Ipsi: $F(1, 18) = 13.56, P=0.0017^{**}$ | Bonferroni's multiple comparisons test<br>0, Contra vs Ipsi: $p>0.9999$<br>1wk, Contra vs Ipsi: $p<0.0001^{****}$<br>2wk, Contra vs Ipsi: $p=0.0054^{**}$<br>3wk, Contra vs Ipsi: $p=0.0121^{*}$<br>4wk, Contra vs Ipsi: $p>0.9999$<br>5wk, Contra vs Ipsi: $p>0.9999$<br>6wk, Contra vs Ipsi: $p>0.9999$<br>7wk, Contra vs Ipsi: $p>0.9999$<br>8wk, Contra vs Ipsi: $p>0.9999$ | C57Bl6: 10 |
| 5-1C | Paired t test:<br>Saline vs 6-OHDA: $t=7.199, df=4, p=0.0020^{**}$ | | Saline: 5<br>6-OHDA: 5 |

#### von Frey assay

Mice were placed under ventilated plexiglass boxes on a wire mesh platform and habituated for at least 2 hours per day for at least 2 days prior to the experiment. On the test day, mice were habituated for at least 30 minutes before the assay. A series of von Frey filaments (North Coast Medical NC12775-02 to NC12775-09) were applied perpendicularly to the point of bending to either the hind paw sural nerve-innervated plantar skin or the hind paw tibial nerve-innervated plantar hairy skin located between the footpads. The nominal bending forces of the filaments, provided by the manufacturer, were 0.02g, 0.04g, 0.07g, 0.16g, 0.4g, 0.6g, 1g, and 1.4g. Paw withdrawal or flinching immediately upon filament application was defined as a positive response. For each given force, the filament was applied 5 times to the ipsilateral (i.e. on side of injury) hind paw. In some cases, the filament was applied 5 times to the contralateral hind paw, then applied 5 times to the ipsilateral hind paw. Intervals between each application were at least a few seconds to avoid sensitization. The number of positive responses out of 5 total applications was used to calculate a given animal’s response frequency. ***Pinprick assay.*** This assay was generally performed at least 30 min after the von Frey assay was completed, on the same platform. An Austerlitz insect pin (000, Fine Scientific Tools) was applied to the hind paw plantar hairy skin. Paw withdrawal or flinching immediately upon filament application was defined as a positive response. Only the ipsilateral hind paw was tested. Intervals between each application were at least a few seconds to avoid sensitization. The number of positive responses out of 5 total applications was used to calculate a given animal’s response frequency. ***Optogenetics.*** Mice were placed under an inverted 500 mL beaker on a glass platform and habituated for at least 2 hours per day for at least 2 days prior to the actual experiment. On the test day, mice were habituated for at least 30 minutes before the assay. A 473 nm Blue DPSS laser (Laserglow technologies, Canada) was used to optogenetically stimulate nerve fibers. 10-100ms light pulses were applied precisely to the tibial area of the plantar surface of the hind paw using an optic fiber cable (Ø400 μm core, M82L01, Thorlabs, Inc., USA). Intervals between each application were at least a few seconds to avoid sensitization. Cre-negative and Cre-positive sham mice were used as controls. The number of positive responses out of 5 total applications was used to calculate a given animal’s response frequency.

### 6-hydroxydopamine injection

For systemic injection, mice received intraperitoneal injections of 6-OHDA (100 mg/kg, Santa Cruz) in saline for 5 consecutive days either at 8 weeks after SNI, or without prior injury. Control mice received intraperitoneal injections of saline. For local injection, under deep isoflurane anesthesia, mice received an injection of 6-OHDA (2μl, 100μg/μl) or saline, delivered via a Hamilton syringe connected to a 30G needle, into the hind paw plantar hairy skin for 5 days.

### Fast Blue Retrograde Labeling

Mice were anesthetized using isoflurane and 1-2μl of FB (1% in saline; Polysciences Inc.) was injected subcutaneously using a Hamilton syringe connected to a 30-gauge needle to label sensory afferents that innervate the injection sites. Retrogradely labeled sensory neurons in DRGs were examined 7 days after injection. To evaluate the subtypes of neurons exhibiting collateral sprouting, FB was injected at the time of SNI surgery or at 7 weeks postoperatively.

### 4-hydroxytamoxifen Injections

4-hydroxytamoxifen (4-HT, Sigma-Aldrich) was dissolved in 100% ethanol (10 mg/ml), mixed with wheat germ oil (Jedwards International Inc., USA), vortexed for 1 min and centrifuged under vacuum for 20-30 min to remove the ethanol. For TH^2ACreER^ animals, 100μl (1mg) of the 4-HT solution was delivered via oral gavage at P13, P14 and P15. For TrkC^CreER^ animals, 20μl (0.1mg) of the 4-HT solution was delivered via intraperitoneal injection at P5.

### Immunohistochemistry

Mice were transcardially perfused with phosphate-buffered saline (PBS) followed by 4% paraformaldehyde (PFA) in PBS and L3-L5 dorsal root ganglia (DRG) and hind paw skin were harvested. For transverse tissue section preparation, tissues were post-fixed in 4% PFA at 4°C overnight. Tissues were cryoprotected in 30% sucrose in phosphate buffer at 4°C overnight, embedded in optimal cutting temperature medium (OCT, Tissue-Tek) and stored at -80°C. Tissues were cryostat sectioned at 10μm for DRG and 16μm for hind paw skin, respectively. DRG and skin sections were thaw-mounted onto glass slides, stored at -80°C, and incubated at 30-37°C for 20 minutes immediately prior to staining. Slides sections were washed with 0.1% Triton X-100 in PBS (PBST.1) 3x 10 minutes. Slides sections were then blocked with 0.3% Triton X-100 in PBS containing 10% normal donkey serum for 1 hour at room temperature. Tissues were incubated overnight with primary antibodies against rat anti-K8 (Univ of Iowa/DSHB, 1:100, #Troma-1), rabbit anti-K17 (from Dr. Pierre Coulombe Univ of Michigan, 1:1000), Chicken anti-NFH (Aves Labs, 1:200, #NFH), goat anti-mCherry (Sicgen, 1:500, #AB0040-500), chicken anti-GFP (Aves Labs, 1:400, #GFP-1020), goat anti-GFP (Sicgen, 1:500, #AB0020), rabbit anti CGRP (ImmnuoStar, 1:1000, #24112), Goat anti-GFRα2 (R&D Systems, #AF429), goat anti CGRP (Abcam, 1:500, #AB36001), Biotin-IB4(Sigma, 1:100, #L2140), sheep anti-TH (EMD Millipore, 1:500, #AB1542), rabbit anti-TH (EMD Millipore, 1:500, AB152), and chicken anti-NeuN (Aves Labs, 1:200, #NUN), at room temperature in a humidity chamber. The following day tissues were washed with PBST.1 3x 10 minutes and then incubated for 1-2 hours at room temperature in a humidity chamber with secondary antibodies; donkey anti-goat Cy3 (Jackson ImmunoResearch, 1:500, #705-166-147), goat anti-chicken 546 (Thermo Fisher, #A11040), donkey anti-rabbit 647 (Jackson ImmunoResearch, #711-605-152), goat anti-rat 488 (Jackson ImmunoResearch, # 112-545-003), donkey anti-chicken 488 (Jackson ImmunoResearch, #703-545-155), donkey anti-rat 488 (Jackson ImmunoResearch, #712-545-153), goat anti-chicken 488 (Thermo Fisher, #A11039), goat anti-rat Cy3 (Jackson ImmunoResearch, #112-165-167), donkey anti-chicken Cy3 (Jackson ImmunoResearch, #703-165-155), donkey anti-goat 557 (R&D Systems, #NL001), and streptavidin-Dylight 405 (Thermo Fisher, #21831). Tissues were then washed with PBS 3x 10 minutes. Floating spinal cord sections were rinsed in water or 0.1M phosphate buffer, mounted on slides, and allowed to air dry. Sections were coverslipped using Dako fluorescence mounting medium (Dako, #S3023).

For whole-mount hind paw skin staining, fat and connective tissue were thoroughly removed to facilitate antibody penetration. Tissues were post-fixed in 4% PFA at 4°C overnight and then briefly washed with PBS to remove excess PFA. Tissues were washed with 1% Triton X-100 in PBS (PBST.hi) 10 x 30 minutes for a total of 5 hours. Tissues were then incubated with primary antibodies diluted in blocking solution (75% PBST.hi, 20% DMSO, 5% normal donkey/goat serum) for 3 days at room temperature. Tissues were washed with PBST.hi 10x 30 minutes and then incubated with secondary antibodies diluted in blocking solution for 2 days at room temperature. Tissues were then again washed with PBST.hi 10 x 30 minutes and dehydrated in serial dilutions of MeOH (50%, 80%, 100% MeOH for 5 minutes each, and an additional 100% MeOH for 20 minutes). Finally, tissues were cleared in BABB (1 volume Benzyl Alcohol to 2 parts Benzyl Benzoate) for 30 minutes and mounted onto slides with BABB. All incubations were done on a rotating/rocking platform.

### In vivo electrophysiology

In vivo DRG electrophysiology recording was performed as previously described (Qu and Caterina, 2016). Briefly, Split^Cre^-Rosa26^LSLtdTomato^ mice were deeply anesthetized using chloral hydrate (500mg/kg). A lateral laminectomy was performed to expose the lumbar DRGs, and oxygenated artificial cerebrospinal fluid (ACSF) was applied to the DRGs in the pool formed by stitching the incised skin into a ring. ACSF contained 130 mM NaCl, 3.5 mM KCl, 24 mM NaHCO_3_, 1.25 mM NaH_2_PO_4_, 1.2 mM MgCl_2_, 1.2 mM CaCl_2_, and 10 mM dextrose, was bubbled with 95% O_2_ and 5% CO_2_ and had a pH of 7.4. Extracellular recordings were obtained from individual sensory neurons whose epineurium was removed and labeled with tdTomato, using a polished suction micropipette. Receptive fields (RFs) of neurons innervating the hind paw of the mouse were identified by probing the skin with a blunt glass probe. Mechanical stimuli were delivered to RFs innervated by DRG neurons, with a 0.5 mm diameter tipped probe using force-controlled indenter (Model 300C-I, Aurora Scientific). Conduction velocity (CV) was obtained by electrically stimulating the skin within the RF using two wire electrodes.

### Image analysis

Images were acquired using a confocal microscope (Nikon A1) and analyzed blinded to surgical time point using NIS elements (Nikon). For hind paw skin whole-mount staining, Z-stack images (approximately 150-200μm in total depth) were acquired across the thickness of the skin tissue. The tibial area of the glabrous skin of the hind paw was defined as 1200μm x 1272μm (width x length) starting from a point 500μm away from the medial hairy-glabrous border. The sural area of the glabrous skin in the hind paw was defined as 800μm x 1272μm in width and length beginning from the boundary of the lateral hairy skin. Hairy plantar skin was defined as the plantar skin region containing hair follicles and bounded by paw pads. For calculation of nerve fiber density from CGRP, NF-H, and TH staining, individual optical slices were subjected to background subtraction using the rolling ball correction tool, at a setting of 7.46 microns, a value empirically determined to eliminate a significant fraction of non-nerve fiber signal, while largely preserving fiber morphology. The resulting optical sections were compiled into maximum projection Z-stacks. Images were thresholded using a single value applied across all images of a given marker and a binary mask was created to define the regions with signal above threshold. The nerve density fraction was then calculated for each fluorescent channel as (area with signal above threshold) / (total area analyzed). In glabrous skin, the entire area was analyzed. In plantar hairy skin, to avoid counting signal from hair shaft autofluorescence, a binary mask corresponding to the visible hair shafts was manually traced and excluded from both the numerator and denominator areas for nerve density calculations. For quantification of tdTomato signal in TrkC^tdTomato^ and Split^Cre^;Rosa26^LSLtdTomato^ lines, quantification of nerve fiber density was the same with two exceptions. First, due to variability in overall staining intensity between tissue batches, thresholds were set for individual contralateral hind paw images, blinded to timepoint, and then applied to the corresponding ipsilateral images from the same mice. Second, for Split^Cre^;Rosa26^LSLtdTomato^ mice, areas of cross-reacting cells with clear non-neuronal morphology were excluded from both the numerator and denominator, as described above for hair shafts. Figures shown throughout the manuscript were subjected only to threshold adjustment, without rolling ball background correction. The impact of image adjustment steps is illustrated in Extended Data Fig. 1-1. The number of lanceolate or circumferential endings formed by labeled fibers in Split^Cre^;Rosa26^LSL-tdTomato^ or TH^2ACreER^;Rosa26^LSL-EYFP^ mice was counted manually by scrolling through the Z stacks of hairy plantar skin. For hind paw transverse section staining, CGRP and GFRα2 intraepithelial nerve fiber (IENF) density per epidermal length was counted in at least 5 sections. Only the number of fibers crossing the dermal-epidermal boundary were counted, not multiple branches of the same fiber. Values for all parameters were expressed either as fractions within a given skin territory, numbers per unit skin area, or numbers per unit skin length, with each symbol shown in figures derived from an individual mouse. For DRG staining, neuronal cell type specific markers were counted as a control for total number of neurons. Approximately 900 neurons per mouse, derived from multiple sections were counted for each data point. The number of FB positive or FB/markers double-positive cells was counted in a blinded manner.

### Experimental Design and Statistical analysis

Full statistical results, including numbers of animals used in each experiment, are included in Table 1. Anatomical studies on C57BL6 mice and behavioral studies were performed on male mice. Anatomical studies on genetically labeled mice were performed on mice of both sexes. For immunostaining, when comparing only two groups, two-tailed Student’s t-test was used for analysis. When comparing only the ipsilateral hind paw across multiple time points, one-way ANOVA was used. When comparing ipsilateral vs. contralateral hind paw across multiple time points, two-way ANOVA was used. For von Frey and pinprick behavioral measurements, when comparing only the ipsilateral hind paw across multiple time points, repeated measures one-way ANOVA was used. When comparing ipsilateral vs. contralateral hind paw across multiple time points, repeated measures two-way ANOVA was used. For optogenetic assays, when comparing genotypes, Mann-Whitney test was used for analysis. Repeated measures two-way ANOVA was used to analyze the effects of group and/or time. ANOVA tests were followed by post-hoc Bonferroni multiple comparisons correction for either multiple times or multiple forces, but not both, in a given comparison. In repeated measures ANOVAs, potential differences related to sphericity were corrected for using the Geisser and Greenhouse method. All data were presented as mean ± SEM and the criterion for statistical significance was p-value<0.05. The exact statistical test used for each experiment and its details can be found in the figure legends and in Table I. In the case of ANOVA analyses, p-values for the overall comparisons between groups are listed on the graphs, and p values at time points derived from the Bonferroni corrections are indicated by asterisks, as defined in the figure legends. All analyses were performed using GraphPad Prism 9.

## Results

### Peripheral nerve injury leads to collateral sprouting of intact sensory nerves

To assay for collateral sprouting of intact nerves into denervated regions after nerve injury, we performed whole mount immunofluorescence (IF) staining of mouse hind paw skin prior to and 1, 4, and 8 wks following spared nerve injury (SNI) surgery (Figure 1A). It has been shown in the rat SNI model that transected nerves do not regenerate past the lesion site (Xie et al., 2017). Thus, any reinnervation in this model is attributable to collateral sprouting. One week after SNI in the mouse, we observed a substantial loss of immuno-reactivity (IR) for peptidergic (CGRP^+^) and myelinated (NF-H^+^) nerve fibers in the middle of the ipsilateral hind paw glabrous skin formerly supplied by the tibial nerve. However, at 4 and 8wks after injury, progressive reinnervation by CGRP^+^ and NF-H^+^ nerve fibers from medial (saphenous) and lateral (sural) territories into denervated glabrous skin was observed (Figure 1B and Extended Data Fig. 1-2). These changes were quantified from 2D projections of confocal image stacks (Figure 1C and 1E), and confirmed our prior qualitative impression of glabrous skin NF-H^+^ fiber loss and restoration after SNI surgery(Jeon et al., 2021).

**Figure 1.**
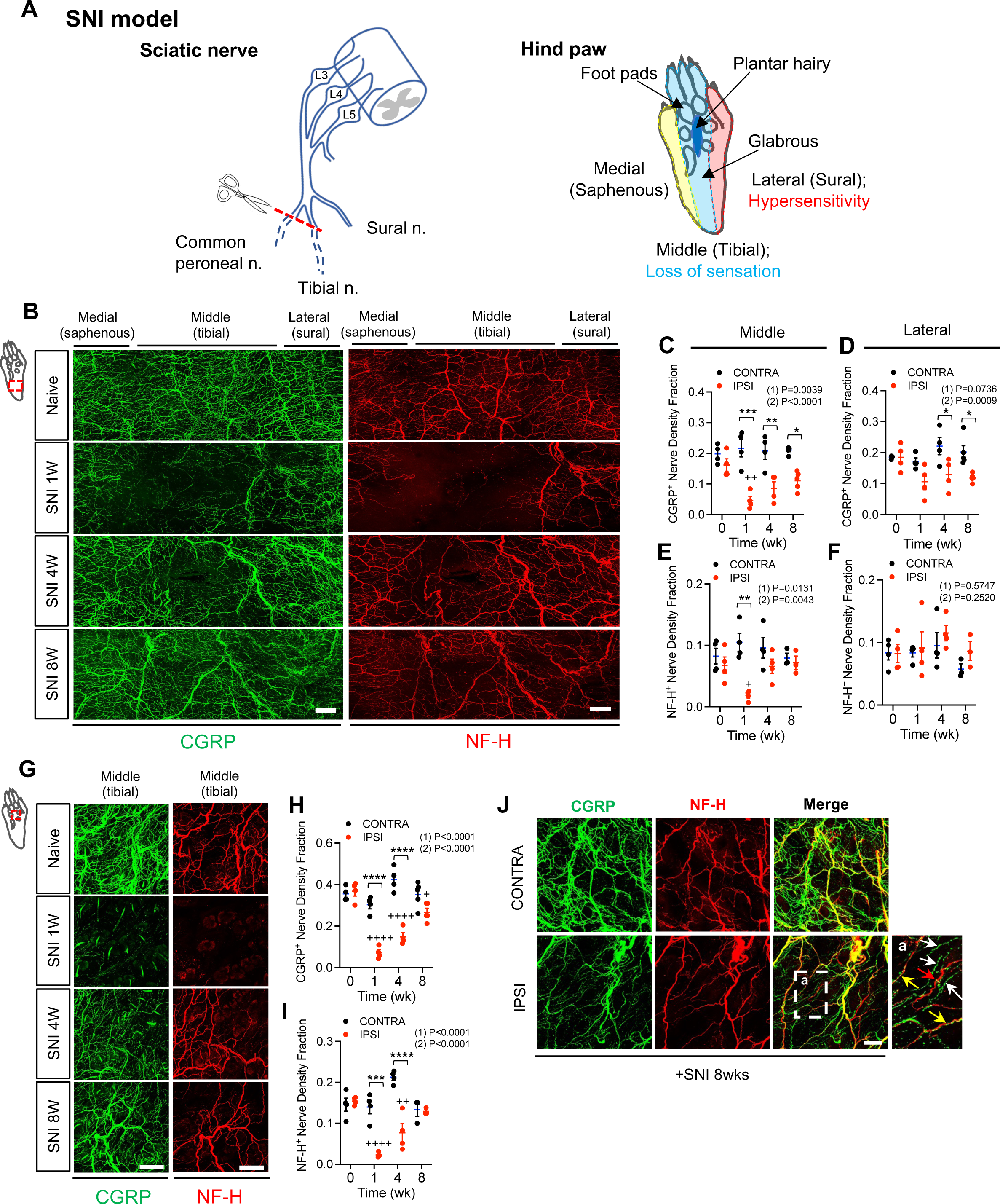
Collateral sprouting of spared nerve fibers after peripheral nerve injury. **(A)** Schematic of the SNI injury model, showing three branches of the sciatic nerve (common peroneal, tibial, and sural) and designating the lateral plantar region (red) of the hind paw innervated by the sural nerve, the middle plantar region of the hind paw innervated by the tibial nerve (blue), and the medial plantar region (yellow) of the hind paw, which is innervated by the saphenous nerve. The dark blue region between the foot pads shows the plantar hairy skin area. **(B)** Whole-mount immunostaining for CGRP (green) and NF-H (red) in the hind paw glabrous skin of C57BL6 mice before and 1, 4, and 8 weeks after SNI surgery. Scale bar, 100μm **(C and E)** Quantification of CGRP (C) and NF-H (E) nerve density fraction in contralateral (black) and ipsilateral (red) middle area of hind paw glabrous skin at the indicated times before and after SNI (n=3-5). **(D and F)** Quantification of CGRP (D) and NF-H (F) nerve density fraction in contralateral (black) and ipsilateral (red) lateral area of hind paw glabrous skin at the indicated times before and after SNI (n=3-5). **(G)** Whole-mount immunostaining for CGRP (green) and NF-H (red) in ipsilateral hind paw plantar hairy skin before and 1, 4, and 8 weeks after SNI surgery. Scale bar, 100μm. Residual green signal at 1 wk is from hair shafts. **(H and I)** Quantification of CGRP (H) and NF-H (I) nerve density fraction in contralateral (black) and ipsilateral (red) plantar hairy skin at the indicated times before and after SNI (n=3-5). **(J)** Double immunostaining for CGRP (green) and NF-H (red) in the hind paw 8 weeks after SNI. Inset (a) shows higher magnification view of the area indicated by dashed box. White arrows indicate CGRP^+^ nerve fibers, red arrows indicate NF-H^+^ nerve fibers, and yellow arrows indicate double-positive fibers, respectively. Scale bar, 100μm. Data are presented as mean ± SEM. In panels C-F, H and I, (1) indicates Overall p-value for difference between baseline and ipsilateral hind paws over time using one-way ANOVA. Results of Bonferroni post hoc correction shown as: ^+^P<0.05; ^++^P<0.01; ^+++^P<0.001; ^++++^P<0.0001. (2) indicates overall p-value for difference between ipsilateral and contralateral paws over time using two-way ANOVA. Results of Bonferroni post hoc correction shown as: *P<0.05; **P<0.01; ***P<0.001; ****P<0.0001.

Next, we quantified CGRP^+^ and NF-H^+^ nerve fibers within a region of the lateral hind paw glabrous skin. This territory is supplied predominantly by the intact sural nerve, but could also have overlapping contributions from the tibial nerve. Indeed, a slight decrease in CGRP^+^ nerve fibers was observed in this lateral hind paw skin territory that persisted until 8 wks after SNI (Figure 1B, 1D, and Extended Data Fig. 1-2). However, no change in sural territory NF-H^+^ nerve fibers was observed at any time point (Figure 1B, 1F, and Extended Data Fig. 1-2). A third skin territory analyzed was that in the vicinity of a group of recently described small hairs between foot pads in the plantar skin of C57BL6 mice (Jeon et al., 2021; Walcher et al., 2018). In this plantar hairy skin, we again saw a significant reduction of CGRP^+^ and NF-H^+^ nerve fibers 1 wk after injury. As in glabrous skin, however, innervation began to return by 4 wks and had nearly fully recovered by 8 wks (Figure 1 G, 1H, 1I and Extended Data Fig. 1-2). Because a subset of CGRP expressing sensory neurons are known to be neurofilament heavy chain-positive, and vice versa, we also examined the overlap between these two markers among reinnervating fibers in glabrous skin. Whereas a subset of fibers were positive for both markers, and are likely to represent myelinated peptidergic nociceptors (Ghitani et al., 2017; Usoskin et al., 2015), CGRP-only and NF-H-only fibers were also observed (Fig. 1J). Some peptidergic nociceptors express a different neuropeptide, Substance P (SP) (Basbaum et al., 2009; Woolf and Ma, 2007). Immunostaining of glabrous and plantar hairy paw skin before and after nerve injury revealed that tibial- (but not sural-) territory SP^+^ fibers are also lost following SNI surgery, but that, like CGRP^+^ fibers, they partially recover by 8 wks (Figure 2A-D). To confirm whether reinnervation of tibial territory skin had arisen by collateral sprouting, rather than regeneration, in a subset of mice, we axotomized the sural nerve 8wks after sural-sparing SNI surgery. While sprouted fibers presumably originating from the more medial saphenous nerve were preserved, there was a loss of fibers on the lateral side of the plantar hairy skin, consistent with a sural nerve origin (Figure 2E). As an independent means of evaluating the subtypes of neurons exhibiting collateral sprouting, we injected a small volume of the retrograde tracer dye, fast blue (FB) into the tibial territory hairy plantar skin of mice at the time of SNI surgery (Fig. 2F). Predictably, compared to the contralateral uninjured side, we observed an ∼4-fold reduction 1 wk after surgery in the proportion of FB labeled cell bodies of lumbar dorsal root ganglion (DRG) neurons. By 8wk after denervation, however, the proportion of retrogradely labeled neurons on the injured side had recovered to approximately 50% of the contralateral control, consistent with these neurons projecting to the previously denervated skin (Fig. 2G, H). Examination of the subtypes of neurons exhibiting retrograde labeling 8wks after injury revealed that recovery was incomplete among CGRP^+^, NF-H^+^, and IB4^+^ (i.e. nonpeptidergic nociceptor/pruriceptor) populations (Fig. 2G, I). There was also a strong trend towards reduction of NFH^+^ but CGRP negative FB+ neurons that was not quite statistically significant (Fig 2I). However, the relative abundance of each marker among those neurons exhibiting apparent sprouting was unchanged from control (Fig 2J).

**Figure 2.**
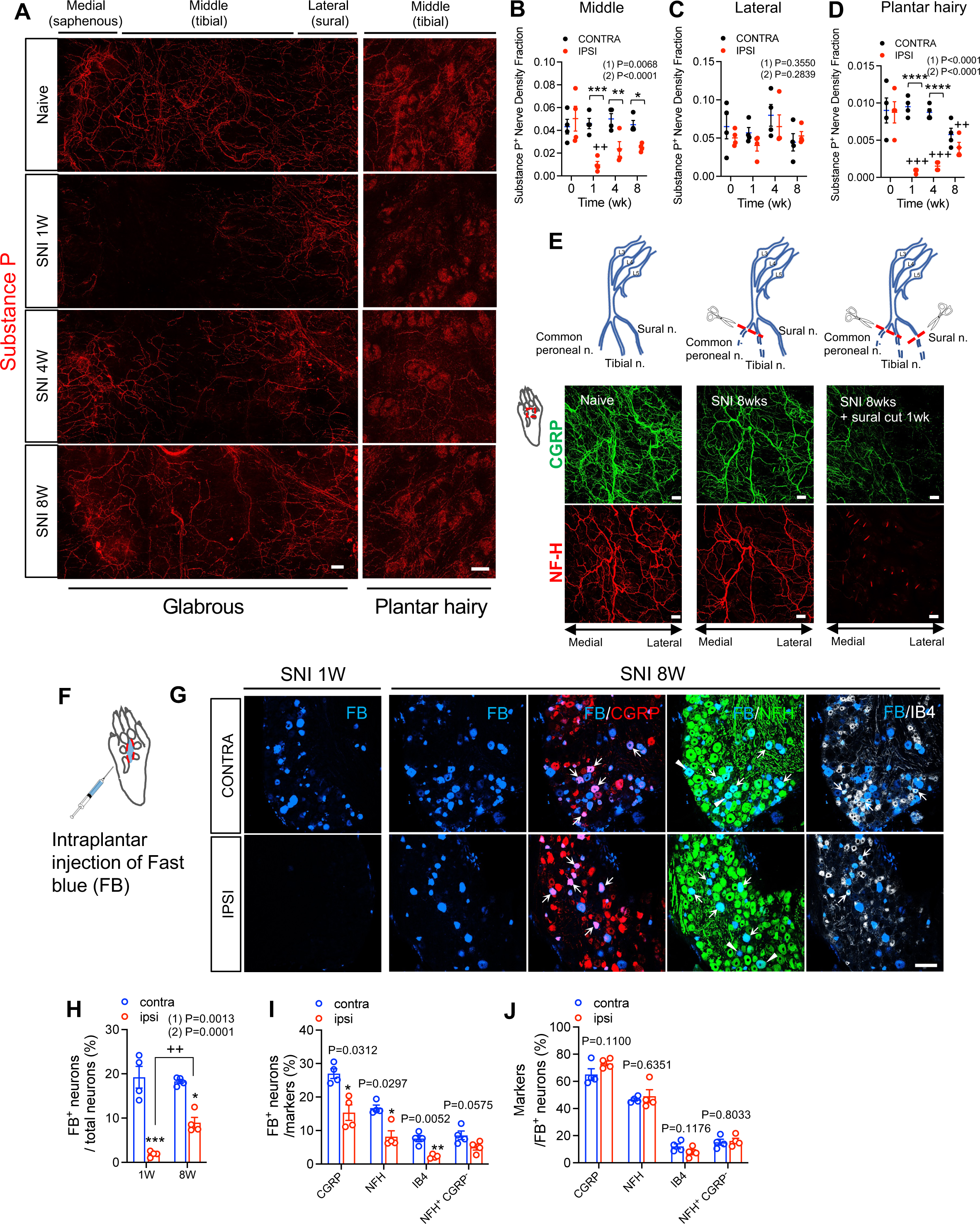
**(A)** Whole-mount immunostaining for Substance P (red) in the hind paw glabrous skin (left panel) and plantar hairy skin (right panel) of C57BL6 mice before and 1, 4, and 8 weeks after SNI surgery. Scale bar, 100μm **(B)** Quantification of Substance P nerve density fraction in contralateral (black) and ipsilateral (red) middle area of hind paw glabrous skin at the indicated times before and after SNI (n=4). **(C)** Quantification of Substance P nerve density fraction in contralateral (black) and ipsilateral (red) lateral area of hind paw glabrous skin at the indicated times before and after SNI (n=4). **(D)** Quantification of Substance P nerve density fraction in contralateral (black) and ipsilateral (red) plantar hairy skin at the indicated times before and after SNI (n=4). **(E)** Schematic showing the site of nerve damage (top panel) either following sural-sparing SNI surgery (middle panel) or sural sparing SNI surgery, followed at 8wks by sural nerve transection (right panel). Bottom panel shows CGRP (green) and NF-H (red) immunostaining in plantar hairy hind paw at indicated time points. Scale bar, 100μm. **(F)** Schematic of intraplantar injection of FB into the hind paw plantar hairy skin. **(G)** FB retrogradely labeled plantar hairy skin-innervating neurons (blue) in the contralateral and ipsilateral lumbar DRGs of C57BL6 mice 1 (left panel) and 8 (right panel) weeks after SNI surgery. Immunostaining for CGRP (red), NF-H(green), and IB4 (white) with FB retrogradely labeled plantar hairy skin-innervating neurons (blue) in lumbar DRGs from C57BL6 mice 8 weeks after SNI surgery. Arrows indicate retrogradely labeled FB/markers doubly positive neurons. Pointed triangles indicate retrogradely labeled FB/NF-H^+^/CGRP^-^ neurons. Scale bar, 100μm. **(H)** Quantification of the percentages of retrogradely labeled FB positive neurons in the contralateral (blue) and ipsilateral (red) lumbar DRGs of C57BL6 mice 1 and 8 weeks after SNI surgery (n=4). **(I)** Quantification of the percentages of FB labeled neurons in CGRP^+^, NF-H^+^, IB4^+^, NF-H^+^ CGRP^-^ neurons in lumbar DRGs from C57BL6 mice 8 weeks after SNI surgery (n=4). **(J)** Quantification of the percentages of CGRP^+^, NF-H^+^, IB4^+^, NF-H^+^ CGRP^-^ neurons in FB labeled neurons in lumbar DRGs from C57BL6 mice 8 weeks after SNI surgery (n=4). Data are presented as mean ± SEM. In panels B-D, (1) indicates Overall p-value for difference between baseline and ipsilateral hind paws over time using one-way ANOVA. Results of Bonferroni post hoc correction shown as: ^+^P<0.05; ^++^P<0.01; ^+++^P<0.001; ^++++^P<0.0001. (2) indicates overall p-value for difference between ipsilateral and contralateral paws over time using two-way ANOVA. Results of Bonferroni post hoc correction shown as: *P<0.05; **P<0.01; ***P<0.001; ****P<0.0001. In panel H, (1) indicates unpaired two-tailed Student’s t-test. ^++^P<0.01, 1wk vs. 8 wk. (2) indicates overall p-value for difference between ipsilateral and contralateral paws over time using two-way ANOVA. Results of Bonferroni post hoc correction shown as: *P<0.05; ***P<0.001. In panels I and J, p-value from paired two-tailed Student’s t-test are shown at top as: *P<0.05, **P<0.01, contra vs. ipsi.

The analyses described above were based upon overall quantification of collateral sprouting into tibial territory skin, without consideration of the skin layers involved. We therefore next assessed intraepithelial nerve fiber loss and reinnervation by immunostaining transverse cryostat sections from hairy plantar skin for either the nonpeptidergic or peptidergic nociceptor/pruriceptor markers. Because IB4 binding exhibited a nonspecific pattern in skin, we instead used GFRα2 as a marker of nonpeptidergic nociceptor/pruriceptor terminals. We first confirmed that there was a substantial, but not exclusive, overlap between IB4 staining and GFRα2 staining among neuronal cell bodies in lumbar DRG (Figure 3A). We further confirmed that epidermal terminal staining for CGRP and GFRα2 in plantar hairy skin of naïve mice was mostly, but not exclusively, nonoverlapping (Fig. 3B). As expected, 1 wk following SNI surgery, we observed a dramatic decrease in both dermal and epidermal innervation by both CGRP^+^ and GFRα2^+^ fibers. Interestingly, at this time point, the disappearance of GFRα2 epidermal staining was not complete. Whether this represents incomplete/delayed degeneration of the transected nonpeptidergic fibers or a minor contribution of innervation by nonpeptidergic fibers of non-tibial (e.g. sural or saphenous) origin is unclear. Nevertheless, we observed evidence of reinnervation by both populations, albeit with distinct kinetics. Reinnervation of the epidermis by CGRP^+^ fibers proceeded gradually and steadily from 1 wk to 8 wks post-injury (Fig. 2 C-E), but did not reach baseline levels in that timeframe. By contrast, reinnervation of epidermis by GFRα2^+^ fibers was profound at 4wks post-injury, achieving levels at or near baseline, but exhibited a trend towards regression between 4wks and 8 wks (Fig. 2 C, F, G). These findings suggest that the forces driving and maintaining sprouting and epidermal penetration by peptidergic vs. nonpeptidergic nociceptive/pruriceptive fibers following nerve injury are distinct.

**Figure 3.**
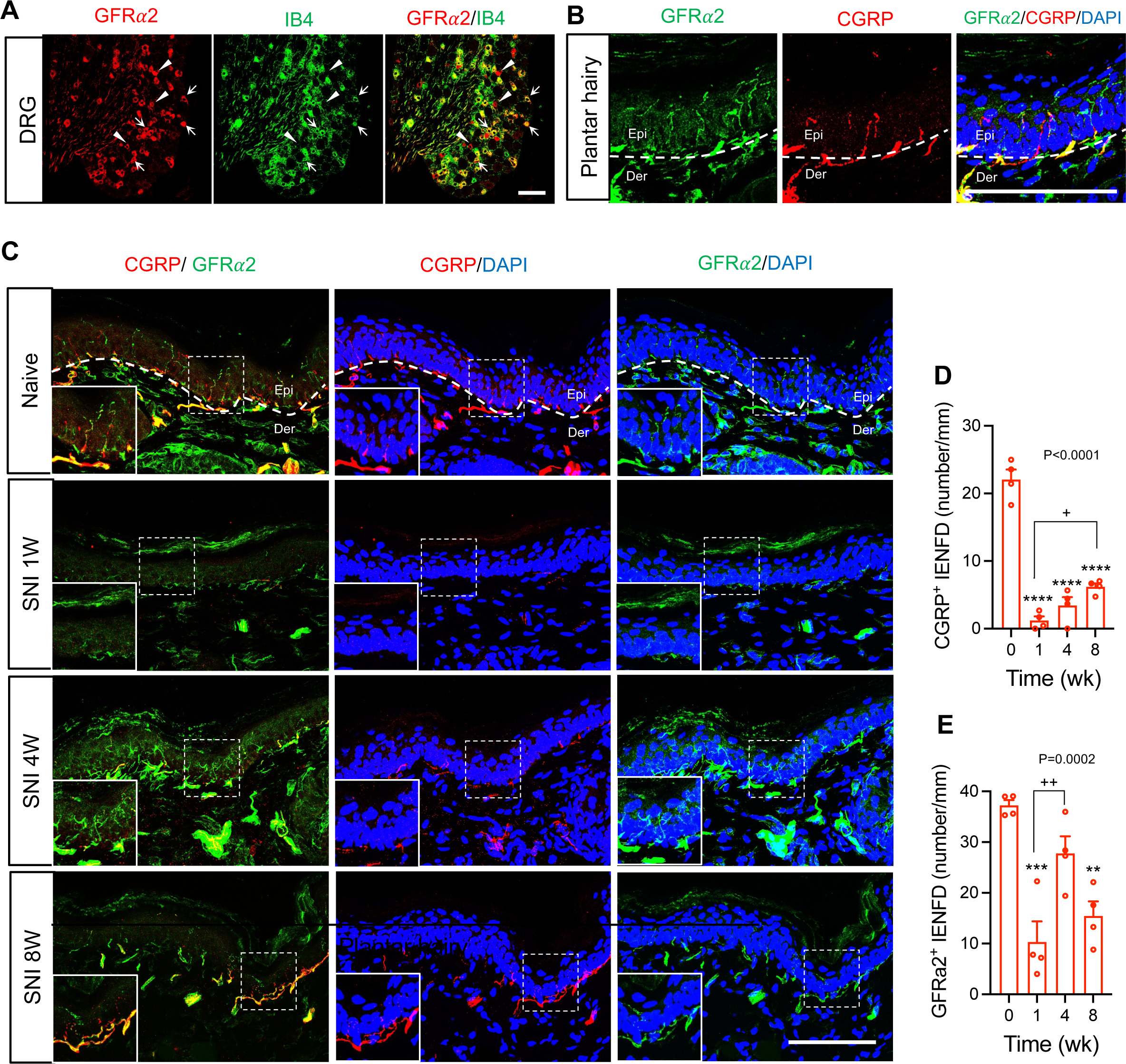
**(A)** Double immunostaining for GFRα2 (red) and IB4 (green) in lumbar DRGs from naïve C57BL6 mice. Arrows indicate colocalization GFRα2/IB4 doubly positive neurons. Arrows indicate neurons showing colocalization of GFRα2 and IB4. Pointed triangles indicate GFRα2 neurons not colocalizing with IB4. Scale bar, 100μm. **(B)** Transverse section double immunostaining for GFRα2 (green) and CGRP (red) in the hind paw hairy plantar skin from naïve C57BL6 mice. White dash is dermal-epidermal boundary. Scale bar, 100μm. **(C)** Transverse section double immunostaining for CGRP (red) and GFRα2 (green) in the ipsilateral hind paw hairy plantar skin before and 1, 4, and 8 weeks after SNI surgery. Curved white dash is dermal-epidermal boundary. Square insets show magnified view of CGRP-IENF and GFRα2-IENF. Scale bar, 100μm. **(D and E)** Quantification of CGRP (D) and GFRα2 (E)-IENF density in the ipsilateral plantar hairy skin at the indicated times before and after SNI (n=4). Data are presented as mean ± SEM. In panel D and E, overall p-value is from one-way ANOVA of ipsilateral paw data over time. Results of Bonferroni post hoc correction shown as: **P<0.01, ***P<0.001, ****P<0.0001, baseline vs. ipsi; ^+^P<0.05, ^++^P<0.01, 1 wk vs. 4wks or 8wks.

### Differential collateral sprouting by Aβ Rapidly Adapting LTMRs vs. TrkC lineage neurons

We next turned to genetically labeled mouse lines to assess the potential of myelinated low-threshold mechanoreceptors (LTMRs), which also express neurofilament heavy chain, to exhibit collateral sprouting. The initial report of hair follicles in mouse hairy plantar skin described their innervation by Aδ-LTMRs and some circumferential fibers (Walcher 2018). To more completely define the baseline LTMR innervation pattern of these hair follicles, we examined them in transgenic mice with selective labeling of additional LTMR subtypes (Figure 4). We first examined Split^Cre^ Rosa26^LSL-tdTomato^ mice, in which Aβ RA-LTMRs are reportedly labeled (Rutlin et al., 2014). In naïve mice of this genotype, we observed tdTomato^+^ fibers and lanceolate endings surrounding hair follicles in plantar hairy skin (Fig. 4A, C). We also observed retrograde labeling of tdTomato^+^ cell bodies by FB injection into the plantar paw skin (Fig. 4A). Because limited physiological validation of the specificity of the Split^Cre^ Rosa26^LSL-tdTomato^ line has been reported, we conducted in vivo electrophysiological recording from the cell bodies of neurons labeled in these mice during controlled mechanical stimulation of the paw. We recorded from 10 tdTomato^+^ neurons in three mice. All 10 exhibited a conduction velocity in the Aβ range. Moreover, in the 4 neurons with mechanical receptive fields in the plantar paw skin, we observed a rapidly adapting response phenotype, including on-and off-responses, in all 4, consistent with their identity as Aβ-RA-LTMRs (Fig. 4B). We next examined TrkC^tdTomato^ mice, in which Aβ SA1-LTMRs and Aβ Field-LTMRs are labeled, in addition to other, less-well described populations (Bai et al., 2015; Rutlin et al., 2014; Li et al., 2011). As we previously reported (Jeon et al., 2021) we observed extensive innervation of hairy plantar skin with TrkC-lineage neurons that include Aβ SA1-LTMRs (Figure 4F, arrow in inset a), defined by their association with Keratin 8 (K8)^+^ Merkel cells, and Aβ Field-LTMRs (Figure 4F, inset b), defined by their circumferential endings. Thus, multiple Aβ-LTMR subtypes innervate plantar hairy skin under basal conditions. We also examined TrkB^CreER^-Rosa26^LSLtdTomato^ mice which, when treated postnatally with 4-hydroxytamoxifen, harbor labeled Aδ-LTMRs (Rutlin et al., 2014). However, while we observed cutaneous afferents in the hairy plantar hind paw skin of these mice, extensive non-neuronal skin cell labeling (data not shown) precluded their effective use for subsequent experiments.

**Figure 4.**
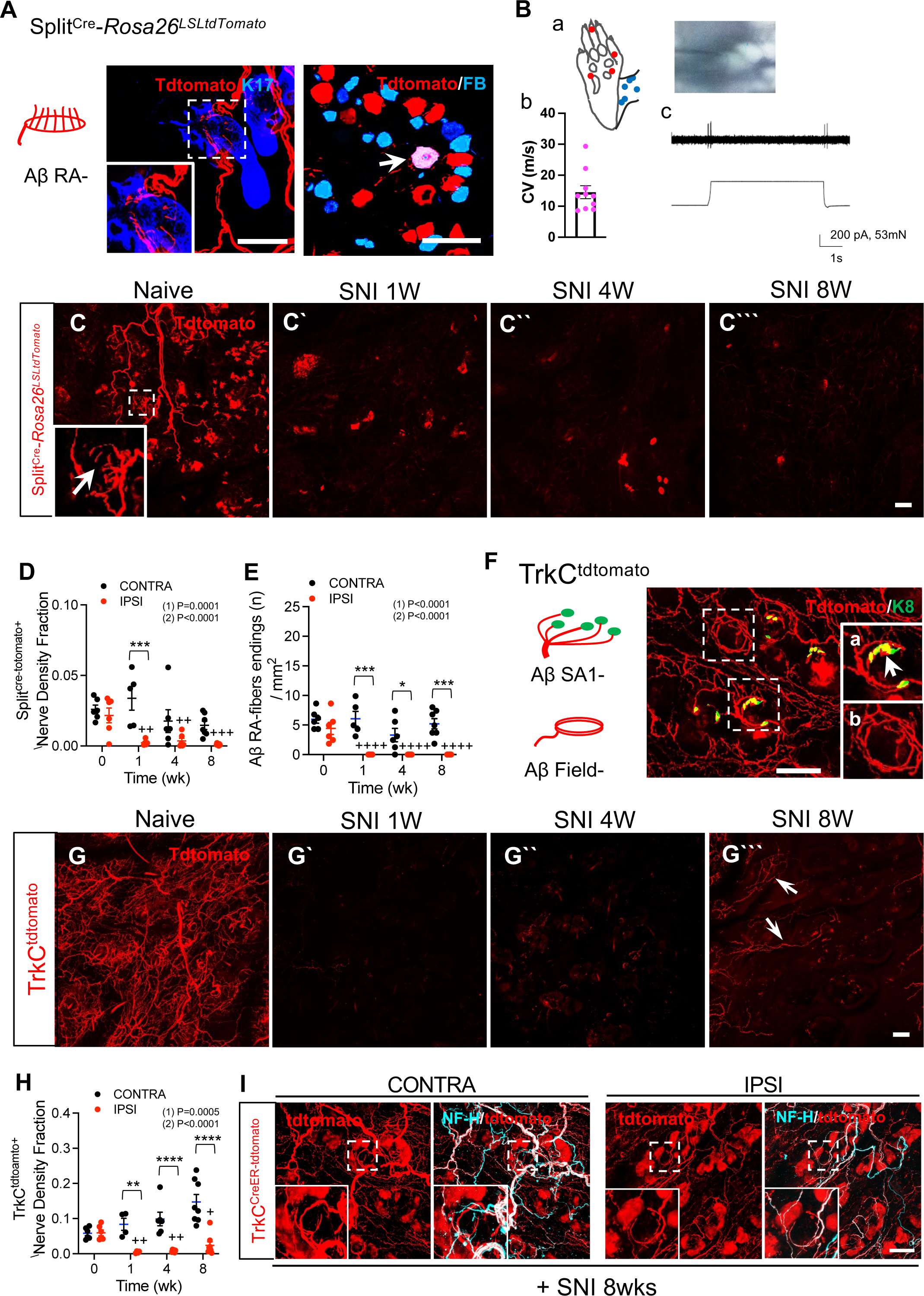
Aβ-LTMRs exhibit absent or limited sprouting into denervated skin after peripheral nerve injury. **(A)** Schematic of characteristic lanceolate endings of Aβ RA-LTMR labeled in Split^Cre^-Rosa26^LSLtdTomato^ mice (left). Modified from Abraira VE et al., Neuron. 2013 Aug 21;79(4):618-39. Whole-mount immunostaining of plantar hairy skin from Split^Cre^-Rosa26^LSLtdTomato^ mice for tdTomato (red) and K17 (blue, hair follicle marker). Arrow in inset indicates lanceolate nerve ending (middle). Immunostaining for tdTomato (red) in FB retrogradely labeled plantar hairy skin-innervating neurons (blue) in lumbar DRGs from Split^Cre^-Rosa26^LSLtdTomato^. Arrow indicates retrogradely labeled FB/tdTomato doubly positive neurons (right). Scale bar, 100μm. **(B)** In vivo electrophysiological recordings in the hind paw from Split^Cre^-Rosa26^LSLtdTomato^ mice (n=4). (a) Schematic showing receptive fields (RF) for all 10 individual neurons used for recording. Red dots represent neurons for which both mechanically stimulation response and conduction velocity (CV) were measured, and blue dots represent neurons for which only CV was measured. (b) A quantitative graph of CV of tdtomato^+^ neurons. (c) Representative in vivo electrophysiological recordings from an Aβ RA-LTMR in response to 80mN mechanical stimulation with a 0.5 mm diameter tipped probe. **(C-C’’’)** In Split^Cre^-Rosa26^LSLtdTomato^ mice, whole mount immunostaining for tdTomato (red) in the ipsilateral plantar hairy skin of the hind paw before (C) and 1(C’), 4(C’’), and 8(C’’’) weeks after SNI. Arrow in inset indicates lanceolate nerve ending. Scale bar, 100μm. **(D)** Quantification of Split^Cre^ Rosa26^LSL-tdTomato^ (n=5-7) nerve density fraction in the contralateral (black) and ipsilateral (red) hind paw plantar hairy skin at the indicated times after SNI. **(E)** Quantification of Split^Cre^-Rosa26^LSLtdTomato^ (Aβ RA-fiber, n=5-7) tdTomato^+^ nerve fiber ending density in the contralateral (black) and ipsilateral (red) plantar hairy skin of hind paw at the indicated times after SNI. **(F)** Whole-mount immunostaining of plantar hairy skin from TrkC^tdTomato^ mice for tdTomato (red) and K8 (green, Merkel cell). Insets show higher magnification views of the areas indicated by dashed boxes. Arrow in inset (a) indicates K8^+^ Merkel cells cluster and associated Aβ SA1 LTMR ending in plantar hairy skin. Inset (b) shows TrkC^tdTomato^ circumferential nerve ending, presumably a field LTMR. Scale bar, 100μm. Images of tissues used in Figure 2F have been presented previously (Jeon et al. 2021). **(G-G’’’)** In TrkC^tdTomato^ mice, whole mount immunostaining for tdTomato (red) in the ipsilateral plantar hairy skin of the hind paw before (G) and 1(G’), 4(G’’), and 8(G’’’) weeks after SNI. Arrows indicate TrkC^tdTomato+^ fibers. Scale bar, 100μm. **(H)** Quantification of TrkC^tdTomato^ (n=5-8) nerve density fraction in the contralateral (black) and ipsilateral (red) hind paw plantar hairy skin at the indicated times after SNI. **(I)** In TrkC^CreER^-Rosa26^LSL-EYFP^ mice with treated with 0.1mg 4-HT at P5, whole mount immunostaining for tdTomato (red) in the ipsilateral plantar hairy skin of the hind paw at 8 weeks after SNI. Insets indicate TrkC^tdTomato+^ fibers. Scale bar, 100μm. Data are presented as mean ± SEM. In panels D,E, and H, (1) indicates overall p-value for difference between baseline and ipsilateral hind paws over time using one-way ANOVA. Results of Bonferroni post hoc correction shown as: ^+^P<0.05; ^++^P<0.01; ^+++^P<0.001; ^++++^P<0.0001. (2) indicates overall p-value for difference between ipsilateral and contralateral paws over time using two-way ANOVA. Results of Bonferroni post hoc correction shown as: *P<0.05; **P<0.01; ***P<0.001; ****P<0.0001.

We then asked whether Aβ LTMR subtypes participate in collateral sprouting after nerve injury, by performing SNI surgery on labeled LTMR mouse lines. One wk after injury, we observed an obvious reduction in nerve fibers in the plantar hairy skin of Split^Cre^ Rosa26^LSL-tdTomato^ (Fig. 4C’, 4D) and TrkC^tdtomato^ (Fig. 4G’, 4H) lines, compared to the contralateral paw (CONT). We also observed a reduction in lanceolate ending structures of Aβ RA-LTMR fibers in Split^Cre^ Rosa26^LSL-tdTomato^ mice (Fig. 4C’, 4E). Endings of Aβ SAI-LTMRs and Aβ-Field LTMRs, labeled in the TrkC^tdtomato^ mice, were not quantified but also appeared to be eliminated (Figure 4G’). In contrast to our findings with nociceptive neurons, however, little or no convincing collateral sprouting of Aβ RA-LTMR fibers into the denervated region or formation of mature endings of these fibers was evident, even 8 wks after SNI (Figure 4C’’’, 4D, E, Extended Data Fig. 4-1). There was some residual tdTomato fluorescence 8wks after injury, but that signal resembled neither nerve fibers nor hair follicles. These results suggest that unlike spared nociceptive neurons, spared Aβ RA-LTMRs fail to sprout into denervated skin after peripheral nerve injury, although we cannot definitively rule out trace amounts of sprouting. In TrkC^tdTomato^ mice, by comparison, we observed heterogeneous results at 8 wks. In most mice, we observed a limited amount of tdTomato^+^ innervation. While such staining was consistently less than that seen in the contralateral paw (Figure 4G’’’ and Extended Data Fig. 4-2), some tdTomato^+^ fibers were evident in most mice. In one mouse (Extended Data Fig. 4-2, mouse #3), sprouting of TrkC positive fibers was robust, and included formation of circumferential endings as well as apparent innervation of sweat glands, although whether the latter was neuronal staining vs. background was less clear. The existence of circumferential endings in the one animal suggests that they include Aβ-Field LTMRs. As a further confirmation of the sprouting of these fibers, we examined TrkC^CreER^ Rosa26^LSL-tdTomato^ mice which were injected postnatally with 4OHT. Prior studies have shown that postnatal activation of Cre in these mice results in labeling of fibers that include Aβ-Field LTMRs (Bai et al., 2015). At 8 wks post SNI, tdTomato^+^ fibers were observed in plantar hairy skin of both contralateral and ipsilateral paws, including apparent circumferential endings that co-labeled with NF-H (Fig. 4I). Together, these data suggest that in contrast to Aβ RA-LTMRs, Aβ-Field LTMRs exhibit collateral sprouting capacity, albeit inconsistently. We also cannot exclude the possibility that some of the sprouting we observed comes from Aβ SAI-LTMRs or from TrkC fibers formerly described to form free endings (Bai et al., 2015), and/or a recently described population of TrkC fibers that innervate the vasculature (Morelli et al., 2021a).

### Collateral sprouting of tyrosine hydroxylase expressing fibers into denervated skin after peripheral nerve injury

Next, we explored whether plantar hairy skin also contains nonmyelinated C-LTMRs, which form lanceolate endings around hair follicles, and can be genetically labeled . in tyrosine hydroxylase (TH)^2ACreER^-Rosa26^LSL-EYFP^ mice by postnatal 4-OHT treatment (Fig. 5A) (Li et al., 2011). In naïve mice of this genotype, we observed nerve fibers positive for enhanced yellow fluorescent protein (EYFP) in plantar hairy skin with endings surrounding hair follicles that exhibited a lanceolate but somewhat disheveled appearance (Figure 5C,F). Both the fibers and their endings colabeled extensively with anti-TH antibody (Figure 5D). A similar pattern of TH labeling was observed in the hairy plantar skin of wild-type C57BL6 mice (Fig. 5G) (Kim et al., 2014). Further evidence for projection of C-LTMRs into hairy plantar skin was obtained via local retrograde FB labeling of neurons innervating this territory in C57BL6 mice, as TH labeling of lumbar DRG neuronal cell bodies revealed a subset of FB/TH double-positive neurons (Fig. 5E). Together, these experiments indicated that at least a subset of the TH positive fibers in naïve hairy plantar skin are of DRG origin and, based on their lanceolate ending structure, are likely C-LTMRs. To examine whether some of the TH^+^ fibers might alternatively arise from sympathetic neurons, we performed intraperitoneal (i.p.) injection of 6-hydroxydopamine (6-OHDA, 100mg/kg) for 5 consecutive days to chemically ablate the sympathetic neurons (Kostrzewa and Jacobowitz, 1974) while sparing C-LTMRs (Li et al., 2011). This treatment failed to eliminate either TH EYFP^+^ or anti-TH^+^ nerve fibers in plantar hairy skin, consistent with the presence of C-LTMRs (Figure 5D’). Interestingly, local intraplantar injection of 6-OHDA eliminated nearly all TH positive nerve fibers (Extended Data Fig. 5-1A, B, C). It is unclear if this represents a non-specific neurotoxicity resulting from a locally administered, high dose of this reagent, as has been reported (Michel and Hefti, 1990), though we observed no overt loss of CGRP^+^ fibers with local 6OHDA treatment (Extended Data Fig. 5-1B).

**Figure 5.**
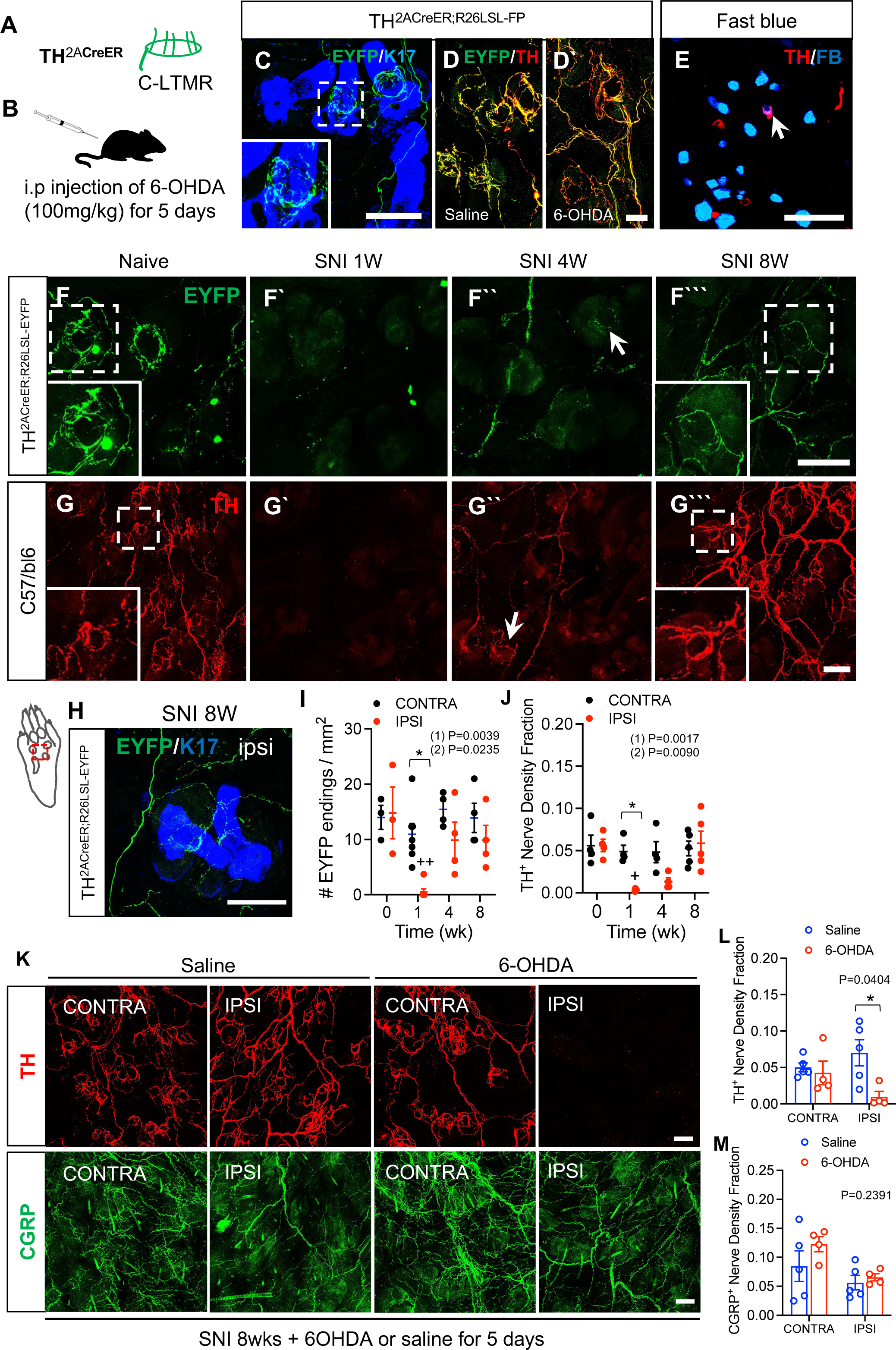
Sprouting of tyrosine hydroxylase expressing fibers into denervated skin after peripheral nerve injury. **(A)** Schematic of characteristic lanceolate endings of C-LTMRs labeled in TH^2ACreER^-Rosa26^LSL-EYFP^ mice. Modified from Abraira VE et al., Neuron. 2013 Aug 21;79(4):618-39. **(B)** Schematic of systemic treatment with 6-OHDA to ablate sympathetic neurons. **(C)** Whole-mount immunostaining for EYFP (green) and K17 (blue) in hind paw plantar hairy skin from TH^2ACreER^-Rosa26^LSL-EYFP^ mice. Inset, expanded view of EYFP^+^ lanceolate nerve ending. **(D)** Whole mount immunostaining for EYFP (green) and TH (red) in the hind paw plantar hairy skin of naïve TH^2ACreER^-Rosa26^LSL-EYFP^ mice after i.p. injection of saline (panel D) and 6-OHDA (panel D’) for 5 consecutive days. **(E)** Immunostaining for TH (red) with FB retrogradely labeled plantar hairy skin-innervating neurons (blue) in lumbar DRGs from TH^2ACreER^-Rosa26^LSL-EYFP^ mice. Arrow indicates retrogradely labeled neuron. Scale bar, 100μm. **(F)** In TH^2ACreER^-Rosa26^LSL-EYFP^ mouse, whole mount immunostaining for tdTomato (red) in the ipsilateral hind paw plantar hairy skin before and 1, 4, and 8 weeks after SNI. Inset, expanded view of GFP^+^ nerve ending. Scale bar, 100μm. **(G)** Whole-mount immunostaining for TH (red) in the ipsilateral hind paw plantar hairy skin of C57BL6 mouse before and 1, 4, and 8 weeks after SNI. **(H)** Whole-mount immunostaining for EYFP+ nerve fibers (green) in TH^2ACreER^-Rosa26^LSL-EYFP^ 8 weeks after SNI, showing relationship of loose circumferential endings to K17+ hair follicles (blue) hair follicles. **(I)** Quantification of EYFP^+^ nerve ending density in contralateral (black) and ipsilateral (red) hind paw plantar hairy skin at the indicated times before and after SNI (n=3-7) in TH^2ACreER^-Rosa26^LSL-EYFP^ mice. **(J)** Quantification of TH^+^ nerve density fraction in contralateral (black) and ipsilateral (red) hind paw plantar hairy at the indicated times before and after SNI (n=4-5) in C56Bl6 mice. **(K)** Whole mount immunostaining for TH (red) and CGRP (green), in the contralateral and ipsilateral hind paw plantar hairy skin of C57BL6 mice 8 wks after SNI and following i.p. injection of saline or 6-OHDA for 5 consecutive days as in panel (B). **(L)** Quantification of TH^+^ nerve density fraction in hind paw hairy plantar skin in the experiment shown in panel (K) (saline, blue, n=4; 6-OHDA, red, n=5). **(M)** Quantification of CGRP^+^ nerve density fractions in the experiment shown in panel (K) (saline, blue, n=5; 6-OHDA, red, n=4). Data are presented as mean ± SEM. In panels I and J, (1) indicates overall p-value for difference between baseline and ipsilateral hind paws over time using one-way ANOVA. Results of Bonferroni post hoc correction shown as: ^+^P<0.05; ^++^P<0.01. (2) indicates overall p-value for difference between ipsilateral and contralateral paws over time using two-way ANOVA. Results of Bonferroni post hoc correction shown as: *P<0.05. In panel L and M, overall p-value for difference between saline and 6-OHDA treated mice using two-way ANOVA. Results of Bonferroni post hoc correction shown as: *P<0.05. Saline vs. 6OHDA.

We next performed SNI surgery on TH^2ACreER^-Rosa26^LSL-EYFP^ and C57BL6 mice to investigate whether intact TH-expressing neurons would sprout into denervated skin after peripheral nerve injury. As with other fiber types examined, we observed a significant reduction in TH EYFP^+^ nerve fiber density 7 days after SNI in TH^2ACreER^-Rosa26^LSL-EYFP^ mice. However, TH EYFP^+^ nerve fibers sprouted new collaterals into denervated skin by 4 wks and 8 wks (Figure 5F-F’’’, 5I). These fibers formed loose circumferential endings around hair follicles (Figure 5F’’’, 5H) that appeared to lack the lanceolate processes characteristic of C-LTMR endings. In C57BL6 mice, we also saw a similar pattern, with significant reduction of TH^+^-IR nerve fibers 7 days after injury and innervation that began to return by 4 wks and that largely recovered by 8 wks (Figure 5G-G’’’, 5J). To determine whether these sprouted fibers included C-LTMRs and/or sympathetic nerve fibers, we injected 6-OHDA intraperitoneally for 5 consecutive days at 8 wks after SNI. In the ipsilateral plantar hairy skin of the 6-OHDA-treated mice, TH^+^ nerve fiber density was significantly reduced compared to that in the saline-treated mice (Figure 5K, L). In contrast, there was no difference in the contralateral plantar hairy skin of the two groups. Systemic treatment with 6-OHDA also did not significantly affect the collateral sprouting of CGRP nerve fibers (Figure 5K, M). Together, these results suggest that whereas many of the TH^+^ nerve fibers observed in control skin are C-LTMRs, TH^+^ fibers exhibiting collateral reinnervation of skin after peripheral nerve injury appear to consist predominantly of sympathetic neurons, without evidence of C-LTMR sprouting.

### Collateral sprouts after nerve injury are functional

To investigate whether the fibers that sprout collaterally after nerve injury are functional, we used optogenetic stimulation of Pirt^Cre^;Rosa26^LSL-ChR2-EYFP^ mice, in which the light-gated ion channel channelrhodopsin-2 (ChR2), fused to EYFP is expressed in the vast majority of DRG neurons. Immunostaining for EYFP confirmed widespread expression not only in nearly all DRG neuronal cell bodies (Figure 6A), but also in their terminals in plantar skin (Figure 6B). A major reduction in ChR2-EYFP^+^ nerve fibers was observed in the hind paw plantar hairy skin 1 wk after SNI. However, intact ChR2-EYFP^+^ neurons sprouted collaterals into denervated skin by 8 wks (Figure 6B). We used a 473 nm laser to optogenetically stimulate these fibers. Light pulses (10 or 100 ms) delivered precisely to the hind paw plantar hairy skin using an optical fiber cable evoked a strong paw withdrawal behavior in naïve Pirt^Cre^;Rosa26^LSL-ChR2-EYFP^ mice, but not controls lacking Cre (Figure 6C). Response frequency to 100 ms blue light stimuli on the ipsilateral hind paw of Pirt^Cre^;Rosa26^LSL-ChR2-EYFP^ mice dropped dramatically by 3 days after SNI, but returned gradually to basal level over 28 days (Figure 6D). The response frequency to the 10 ms blue light stimulus on the ipsilateral hind paw also dropped dramatically by day 3 after SNI and gradually recovered, but did not reach basal level by 28 days (Figure 6E). These findings indicated substantial, but not complete, recovery of functional sensory afferents within this time frame.

**Figure 6.**
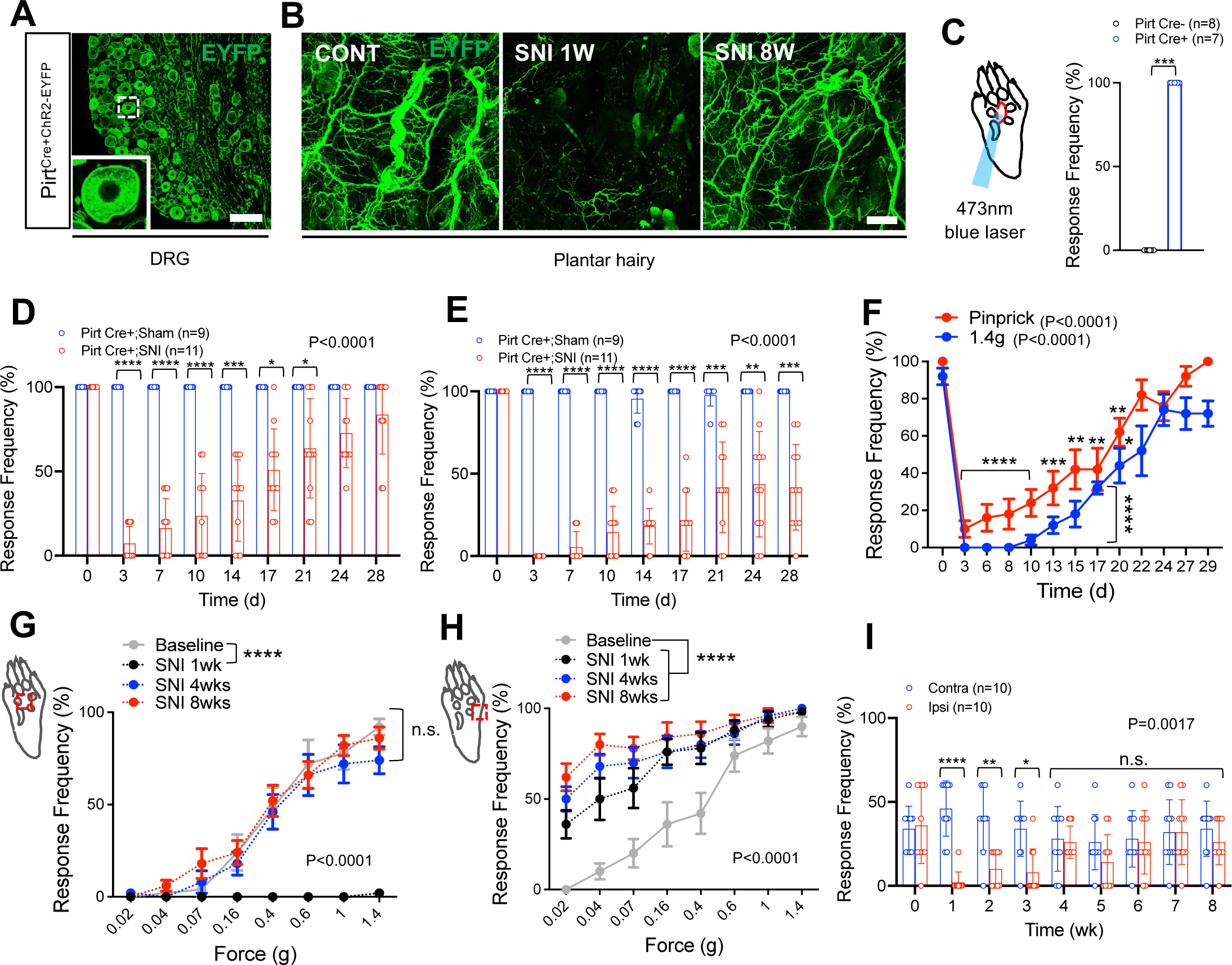
Collateral sprouting contributes to functional recovery of sensory neurons after peripheral nerve injury. **(A)** Immunostaining for EYFP (green) in lumbar DRGs from Pirt^Cre+^;Rosa26^LSL-ChR2-EYFP^ mice. Scale bar, 100μm. **(B)** Whole mount immunostaining for EYFP (green) on the contralateral and ipsilateral hind paw plantar hairy skin before and 1 and 8 weeks after SNI in Pirt^Cre+^;Rosa26^LSL-ChR2-EYFP^ mice. **(C)** Schematic of mouse paw stimulation using 473 nm blue laser (left) and the frequency of withdrawal from light stimulation of the hind paw of Pirt^Cre+^;Rosa26^LSL-ChR2-EYFP^ and Pirt^Cre-^;Rosa26^LSL-ChR2-EYFP^ mice (right). Data are presented as mean ± SEM. Mann-Whitney test. ***P=0.0002, Cre-vs. Cre+. **(D)** Time course of 100 ms blue light-induced paw withdrawal frequency in the ipsilateral tibial nerve-stimulated plantar hair skin after SNI (n=11) and sham (n=9) surgery in Pirt^Cre+^;Rosa26^LSL-ChR2-EYFP^ mice. In panels D, E, and I, circles represent data from individual mice. **(E)** Time course of 10 ms blue light-induced paw withdrawal frequency in the ipsilateral tibial nerve-stimulated plantar hair skin after SNI (n=11) and sham (n=9) surgery in Pirt^Cre+^;Rosa26^LSL-ChR2-EYFP^ mice. In panels **D and E,** two-way repeated measures ANOVA with p-value from overall comparison between groups over time shown at top. Results of Bonferroni post hoc correction shown as: *P<0.05; **P<0.01; ***P<0.001; ****P<0.0001, sham vs. SNI. All quantified data are presented as mean ± SEM. In panels C, F, G, and H, symbols represent individual mice. **(F)** Time course of pinprick (red) or 1.4g von Frey filament (blue) evoked mechanical sensitivity, after SNI, in the ipsilateral tibial nerve-innervated plantar hairy skin of C57BL6 mice (n=10). One-way repeated measures ANOVA with p-value for difference between baseline and ipsilateral hind paws over time shown at bottom. Results of Bonferroni post hoc correction shown as: *P<0.05; **P<0.01; ****P<0.0001. **(G)** Punctate mechanical sensitivity measured across forces in the ipsilateral tibial nerve-innervated plantar hairy skin of C57BL6 mice at baseline (grey), and 1 (black), 4 (blue), and 8 weeks (red) after SNI (n=10). Two-way repeated measures ANOVA with p-value from overall comparison between groups over time shown at bottom. Results of Bonferroni post hoc correction shown as: ****P<0.0001, baseline vs. SNI. **(H)** Punctate mechanical sensitivity measured across forces using von Frey filaments in the ipsilateral sural nerve-innervated lateral hind paw plantar skin at baseline (grey), and 1 (black), 4 (blue), and 8 weeks (red) after SNI (n=10). Two-way ANOVA with p-value from overall comparison between groups over time shown at bottom. Results of Bonferroni post hoc correction shown as: ****P<0.0001, baseline vs. SNI. **(I)** Time course of hind paw sensitivity to 0.4g von Frey filaments, after SNI, in the contralateral and ipsilateral tibial nerve-innervated plantar hairy skin of C57BL6 mice (n=10). Two-way repeated measures ANOVA with p-value from overall comparison between groups over time shown at top. Results of Bonferroni post hoc correction shown as: *P<0.05; **P<0.01; ****P<0.0001, contra vs. ipsi.

Given the altered complement of sensory fibers in the plantar hairy skin after collateral sprouting, namely a predominance of nociceptors and a paucity of LTMRs, we sought to quantitatively examine responses to natural, tactile stimuli. We therefore measured behavioral responsiveness to punctate mechanical stimuli, applied to the plantar hairy skin of the hind paw. Application of either a high force (1.4g) von Frey filament or an Austerlitz pin (Latremoliere et al., 2018) evoked paw withdrawal with high efficiency in naïve C57BL6 mice. Three days after SNI surgery, the response frequencies to both stimuli, when delivered to the denervated hind-paw skin, decreased markedly. However, responsiveness returned gradually to basal levels by ∼4 wks (Figure 6F). Assay with von Frey filaments over a full range of forces confirmed that SNI produced a loss of touch-evoked responses at 1 wk. Yet, touch sensitivity recovered to a pattern indistinguishable from baseline by 4wks, and exhibited a similar pattern at 8wks with no shift in the stimulus-response curve, suggesting there was a lack of mechanical hypersensitivity at either time (Figure 6G). We further confirmed the absence of mechanical hypersensitivity throughout this time window by performing the von Frey assay once a week with a filament of intermediate force (0.4g) applied to the plantar hairy skin (Figure 6I). In contrast, the expected SNI-induced mechanical hypersensitivity was observed in the lateral plantar hind paw skin (sural dermatome) at 1 wk and persisted through 8 wks (Figure 6H). These results suggest that collateral sprouting contributes to functional recovery of sensory neurons in denervated skin, and that even behavioral responses to relatively weak mechanical stimuli can be restored, despite the apparently limited participation of LTMRs in collateral sprouting.

## Discussion

In this study, we assessed collateral sprouting following SNI surgery in the mouse. After initial denervation in tibial nerve skin territory, we observed reinnervation attributable to collateral sprouting of neighboring uninjured afferents. However, sprouting was not uniform among neuronal subtypes. Peptidergic nociceptors, both myelinated and unmyelinated, showed extensive sprouting into denervated skin. Unmyelinated nociceptors/pruriceptors showed a burst of sprouting by 4 wks, but those terminals exhibited a trend towards regression by 8 wks. Aβ RA-LTMRs and C-LTMRs, both of which innervated hairy plantar skin at baseline, failed to convincingly sprout, even 8 wks after denervation. However, a subset of TrkC lineage neurons, including some that were myelinated and most likely Aβ field LTMRs exhibited occasional sprouting.

Prior studies in rat reported collateral sprouting by spared peptidergic and NF-H-positive sensory fibers (Nascimento et al., 2015; Cobianchi et al., 2014; Duraku et al., 2012; Duraku et al., 2013). Another study used the mouse SNI model and serial in vivo microscopy to show that sural nerve nociceptive neurons sprouted collaterally into digit-tip glabrous skin in the tibial nerve territory (Gangadharan et al., 2022). In our study, as well as several performed in rat (Nascimento et al., 2015; Cobianchi et al., 2014; Duraku et al., 2013; Lakatos et al., 2020), sprouting peptidergic and nonpeptidergic neurons reinnervated not only the upper dermis, but also the epidermis. In contrast, Gangadharan et al. reported that sprouting nociceptive neurons remained at the dermal-epidermal boundary, without epidermal penetration, even 42 weeks post-injury (Gangadharan et al., 2022). This difference may be related to the distinct anatomical site (i.e. digit tip) assayed in that study. Further evidence for nociceptor collateral sprouting in rat has come from functional assays (Doucette and Diamond, 1987; Lakatos et al., 2020). In the mouse SNI model, skin-nerve preparation electrophysiological recordings directly showed expansion of receptive fields of sural nociceptive C fibers into tibial skin territory (Gangadharan et al., 2022).

Our finding that some LTMRs show little or no collateral sprouting is generally in line with prior reports using behavioral, anatomical, or electrophysiological methods (Gangadharan et al., 2022; Jackson and Diamond, 1984b). However, as noted above, in a subset of mice we did observe sprouting by TrkC lineage neurons that morphologically resembled Aβ-Field LTMRs. Should this be confirmed electrophysiologically, it might suggest an inefficient LTMR sprouting process potentially amenable to augmentation, if therapeutically warranted.

We also observed innervation of hairy plantar skin by TH^+^ nerve fibers before and 8wks after SNI surgery. In naïve mice, many of these were resistant to ablation by systemic 6-OHDA, consistent with C-LTMRs (Morelli et al., 2021). In contrast, 8 wks after SNI surgery, sprouted TH^+^ afferents (but not TH^+^ fibers in the naïve contralateral paw) were largely eliminated by systemic 6-OHDA, consistent with a predominantly sympathetic neuron origin (Michel and Hefti, 1990). Prior studies have reported sympathetic collateral sprouting into denervated rat skin (Gloster and Diamond, 1995; Nascimento et al., 2015). In one such study (Nascimento et al., 2015), the sympathetic fibers were present only transiently (4-6 wks), whereas in our present study, they persisted at 8 wks. Gangadharan et al. observed cutaneous sprouting by TH^+^ fibers in a subset of denervated mice, but inferred a DRG origin (Gangadharan et al., 2022). Sympathetic neurons also sprout into the DRG following nerve injury and contribute to spontaneous firing by clusters of neurons and to spontaneous pain behaviors (Zheng et al., 2022). Details of sympathetic neuron structure and function in naïve and reinnervated plantar skin therefore merit future attention. We also cannot exclude the presence of TH^+^ TrkC lineage sensory neurons that form endings around blood vessels rather than hair follicles (Morelli et al., 2021b).

Why might collateral sprouting be heterogeneous among neuronal subtypes? One feature shared by sympathetic and peptidergic neurons is responsiveness to nerve growth factor (NGF). Moreover, anti-NGF antibodies suppress collateral sprouting, but not conventional regeneration, by both neuronal populations (Diamond et al., 1987; Diamond and Foerster, 1992; Gloster and Diamond, 1995). However, the lack of NGF responsiveness of adult nonpeptidergic nociceptors, which also sprout, albeit with an apparently transient timecourse, suggests involvement of additional factors whose abundance might exhibit similarly transient kinetics. Other possible contributors to differential sprouting include differential intrinsic activity and/or mechanosensitivity (Gangadharan et al., 2022), differential expression of sprouting associated genes (Harrison et al., 2015; Lemaitre et al., 2020; Gangadharan et al., 2022) or differential expression of receptors for factors released during Wallerian degeneration, which may promote sprouting (Collyer et al., 2014). We also cannot exclude the possibility that the apparent difference in sprouting propensity between Aβ-RA-LTMRs and TrkC lineage fibers reflects the greater baseline cutaneous density of the latter.

In parallel with a sequence of denervation, then gradual reinnervation, we observed a loss and then gradual recovery of behavioral responses to mechanical stimulation in denervated hairy plantar skin that reached baseline levels by 4 wks. Responses to optogenetic activation of sensory neurons were also lost after injury, but partially recovered towards baseline levels within 4 wks. These data confirm the functionality of collaterally sprouted fibers. Lacking a system that would permit us to focus heat specifically on hairy plantar skin, we could not monitor thermosensory recovery. Our functional studies revealed two interesting findings. First, we saw full recovery of von Frey force-response relationship, including at low forces, despite an incomplete repertoire of LTMR sprouting. Prior data have implicated MrgprD^+^ nonpeptidergic nociceptors/pruriceptors, but not peptidergic C fibers, in the response to punctate mechanical stimuli in naïve mice (Cavanaugh 2009). Yet, significant von Frey responses remain even after nociceptor ablation (Abrahamsen 2008). LTMRs, on the other hand, have been proposed to qualitatively shape and in some cases attenuate these responses (Arcourt et al., 2017). More sophisticated behavioral assays, including high-speed video analysis (Arcourt et al., 2017; Abdus-Saboor et al., 2019) will be required to determine whether the sprouted nociceptors recapitulate the full range of pre-injury mechanical responsiveness. Such assays should also be correlated with TrkC lineage labeling, to determine whether inconsistent sprouting by these fibers influences behavioral responses. Second, despite the gradual recovery of normative punctate nociception, at no time during the 8wk course of our experiments did we observe mechanical hypersensitivity in the reinnervated area. In the rat sciatic nerve transection model, Cobianchi et al. reported loss of sensory function in denervated skin, followed by recovery, with a brief, transient period of mechanical hypersensitivity ∼50 days after sciatic transection (Cobianchi et al., 2014). A prior study of humans undergoing C7 spinal nerve transposition surgery reported transient mechanical and cold hypersensitivity in denervated skin in a subset of patients that was likely attributable to collateral sprouting (Ali et al., 2002). Extensive analysis of sensory recovery from denervation following SNI in the mouse was performed by Gangadharan et al. (Gangadharan et al., 2022). Within the time window common between our two studies, our respective findings are similar, namely full and faithful recovery of von Frey force-response characteristics in the tibial territory by 8 wks. Remarkably, however, while in their study mechanosensitivity remained relatively stable for the ensuing 8 wks, 16 wks after SNI the mice began to develop mechanical hypersensitivity in the tibial territory that was paralleled by supranormal sensitivity of C fibers innervating tibial skin. Concomitant with this hypersensitivity, the authors observed *de novo* innervation of Meissner’s corpuscles in tibial territory digit tip skin by nociceptive fibers, providing one possible explanation for their mechanical hypersensitivity. We do not know if hyperalgesia would also have been seen in our mice at times beyond 8wks. It is also not clear to what extent glabrous vs. hairy plantar skin was stimulated during the functional studies of Gangadharan et al, though their skin-nerve recordings appear to have focused predominantly on glabrous skin. It is also possible that the anatomical differences between our two studies, such as presence vs lack of epidermal reinnervation, sprouting by TrkC lineage neurons, or prevalence of sympathetic sprouting would have functional consequences at later time points.

In summary, we provide evidence for subtype-specific collateral reinnervation of hairy plantar skin in the mouse SNI model that is accompanied by robust recovery of punctate mechanosensitivity within 4 wks, despite a paucity of LTMRs. Recovery of tactile sensation was seen without hypersensitivity, despite the fact that a portion of the sprouting originated from the adjacent sural dermatome, which demonstrated characteristic long-lasting hypersensitivity. Collectively, our results highlight the importance of future studies to determine what drives subtype-specific sprouting and how neuronal plasticity varies across the receptive fields of reinnervated and spared skin territories. They also have implications for strategies to enhance sensory recovery and limit neuropathic pain after peripheral nerve injury.

## Acknowledgements

This work was supported by grants from the Blaustein Pain Research Program and Merkin Peripheral Neuropathy and Nerve Regeneration Program at Johns Hopkins (S.M.J.), by the Neurosurgery Pain Research Institute at Johns Hopkins (M.C., S.M.J., A.L.), by R01 NS112266 (A.L.), by the Johns Hopkins Summer Internship Program (Y.C.) and by K12 DK1000022 (L.K.C.) The authors are grateful to Drs. David Ginty (Harvard Univ.) and Xinzhong Dong (Johns Hopkins Univ.) for generously sharing mouse lines, to Pierre Coulombe for sharing anti-K17 antibodies, to John Robinson, Ian Reucroft, and Gabriella Muwanga for assistance with mouse colony management, and to Lintao Qu, Allan Belzberg, and members of our labs for helpful feedback.

## Author contributions

M.J.C. and S.M.J. conceived of the project. M.J.C., A.P., L.K.C., D.C., and S.M.J. designed research. S.M.J. and D.C. performed research, S.M.J., A.P., L. M., C.Y., and L.K.C. analyzed data and all authors participated in writing and/or editing the manuscript.

---

**Figure 1-1.**
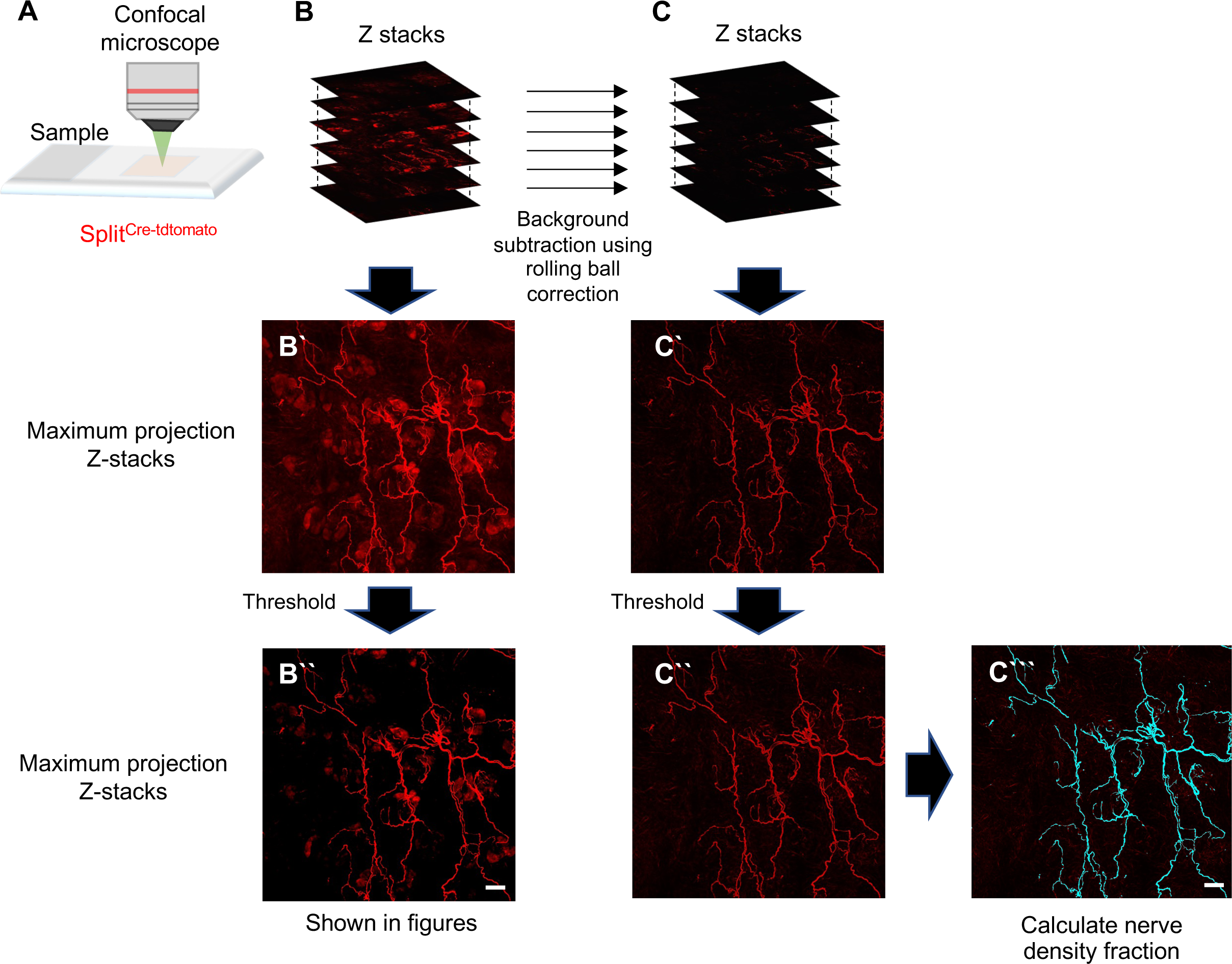
Image processing steps for figure preparation and nerve density fraction calculation. **(A)** Schematic of confocal image acquisition. **(B)** For preparation of most figures, confocal z-stacks images (B) were converted to a maximum intensity 2D projection image (B’) and thresholded to generate a representative image (B’’). Scale bar, 100μm **(C)** For nerve fraction quantification, Z stack images (C) were subjected to background subtraction prior to being converted to 2D maximum intensity projection images (C’). The remaining signal was thresholded (C’’) and signal above threshold converted to a binary image (C’’’), which was then used to calculate nerve density fraction (i.e. fraction of area in which above-threshold signal is detected). Scale bar, 100μm.

**Figure 1-2.**
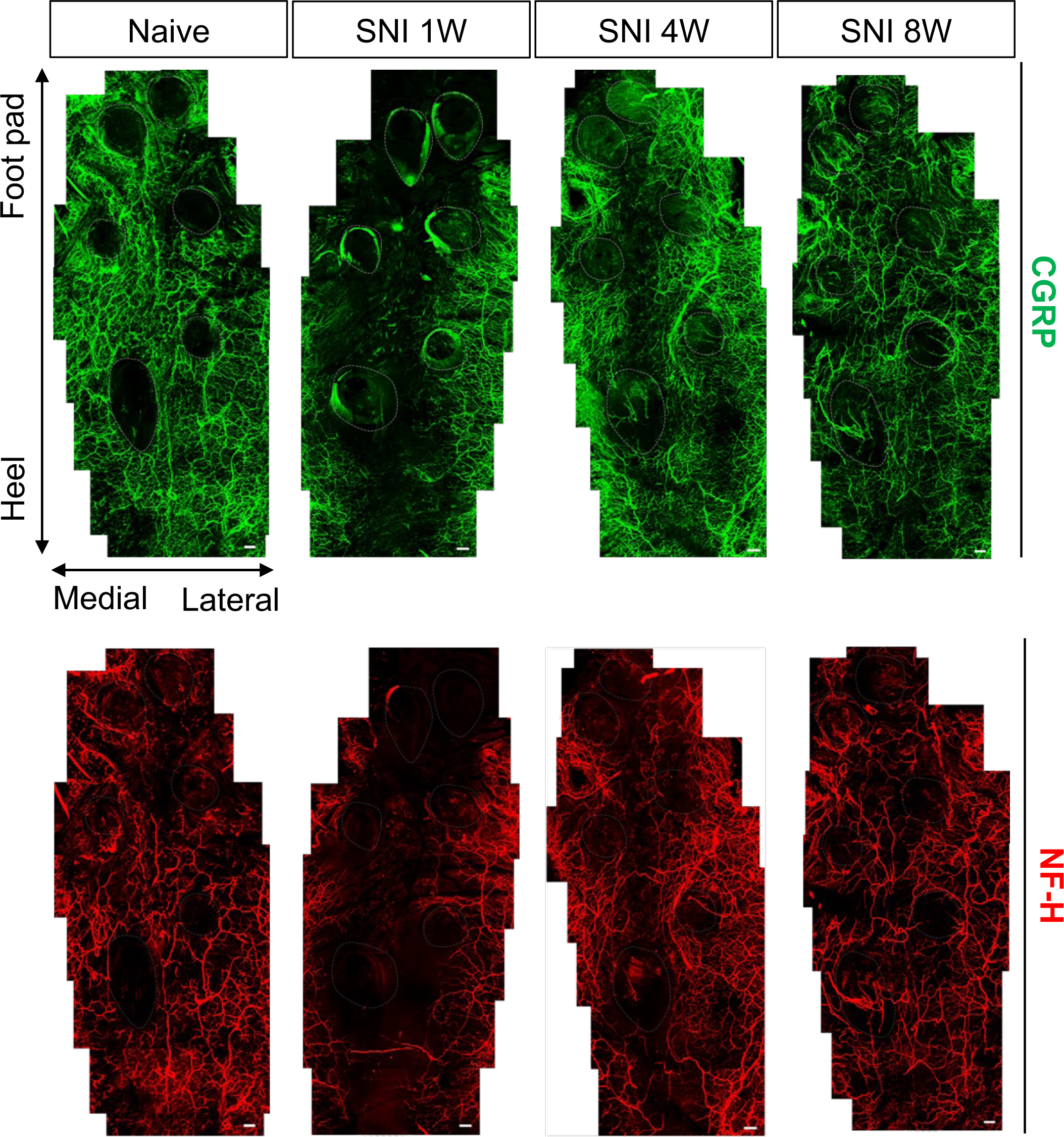
Tiled images of hind paw plantar skin (digits omitted) following whole-mount immunostaining for. CGRP (green) and NF-H (red) before and 1, 4, and 8 weeks after SNI. Footpad skin was not imaged. Scale bar, 100μm

**Figure 4-1.**
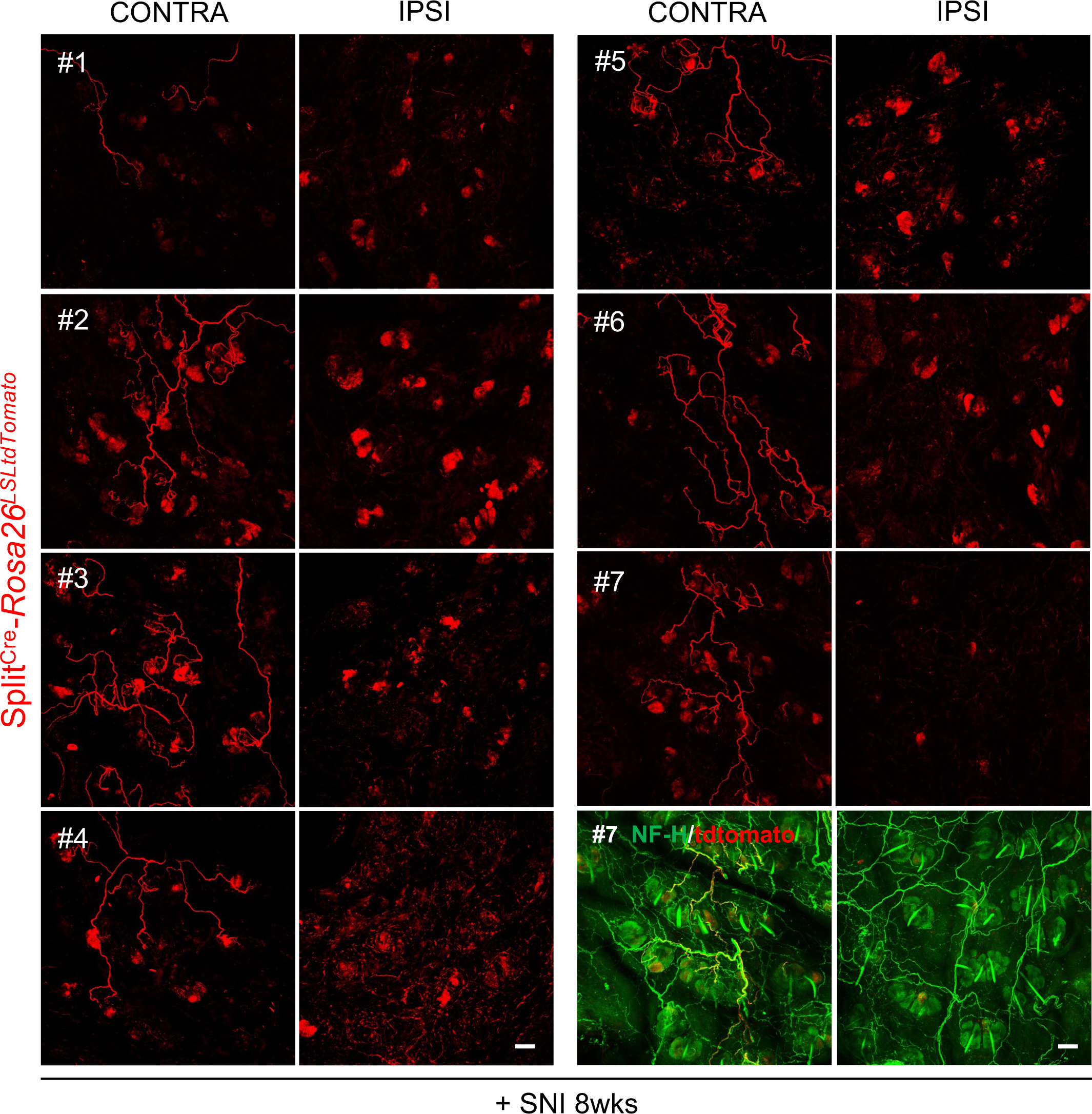
Collateral sprouting of Aβ-RA-LTMRs after peripheral nerve injury. Immunostaining for tdTomato (red) in the contralateral and ipsilateral hind paw plantar hairy skin of individual Split^Cre^-Rosa26^LSLtdTomato^ mice 8 weeks after SNI. The last panels at bottom right show double IF staining for NF-H (green) and tdtomato (red) in the ipsilateral hindpaw plantar hairy skin of mouse #7. Scale bar, 100μm.

**Figure 4-2.**
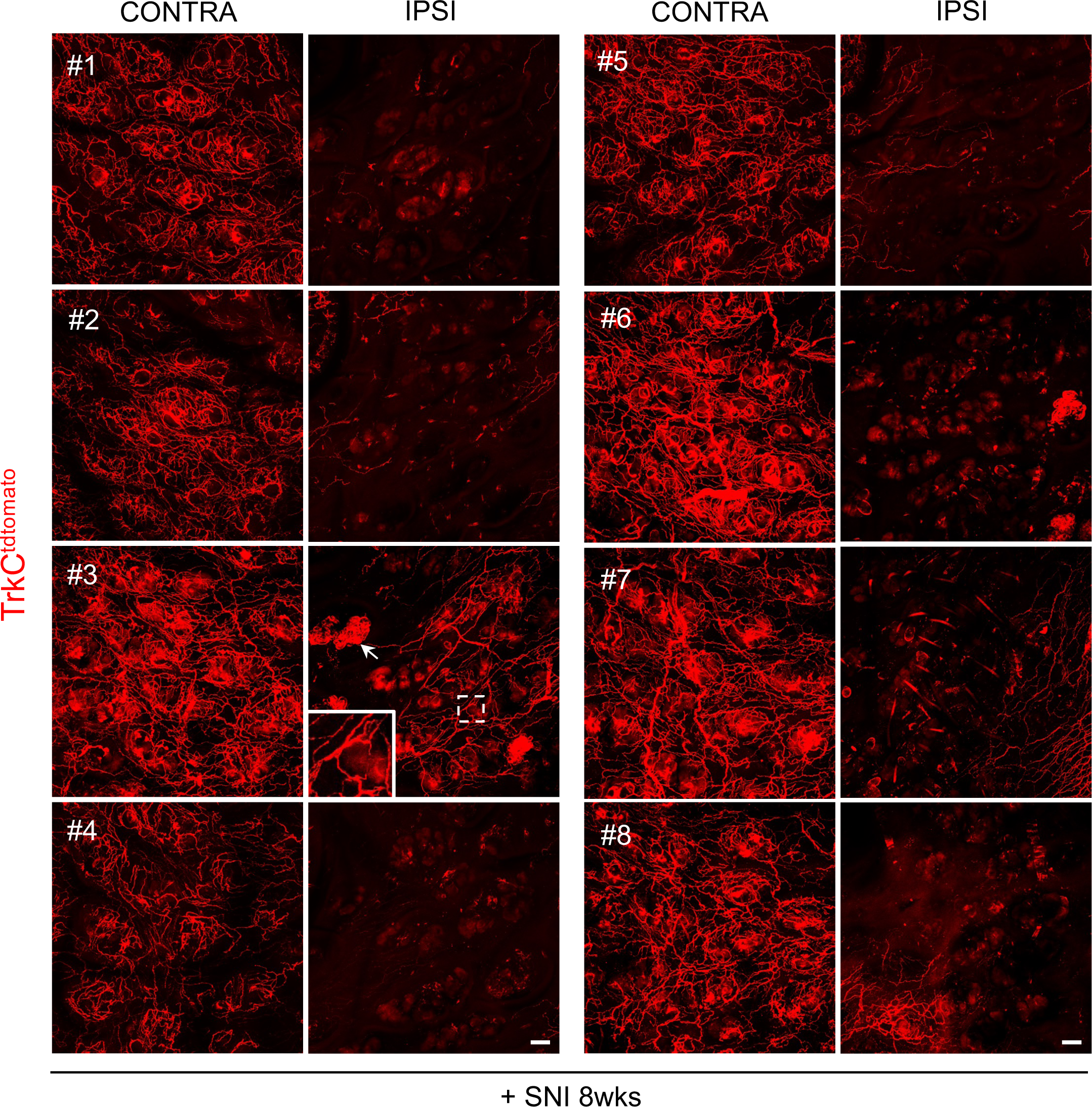
Collateral sprouting of TrkC^tdTomato^ nerve fibers after peripheral nerve injury. Immunostaining of tdTomato (red) in the contralateral and ipsilateral hind paw plantar hairy skin of individual TrkC^tdTomato^ mice 8 weeks after SNI. In the ipsilateral hind paw plantar hairy skin of mouse #3 (right), white arrows indicate possible innervation of sweat glands. Inset shows TrkC^tdTomato^ circumferential nerve ending. Scale bar, 100μm.

**Figure 5-1.**
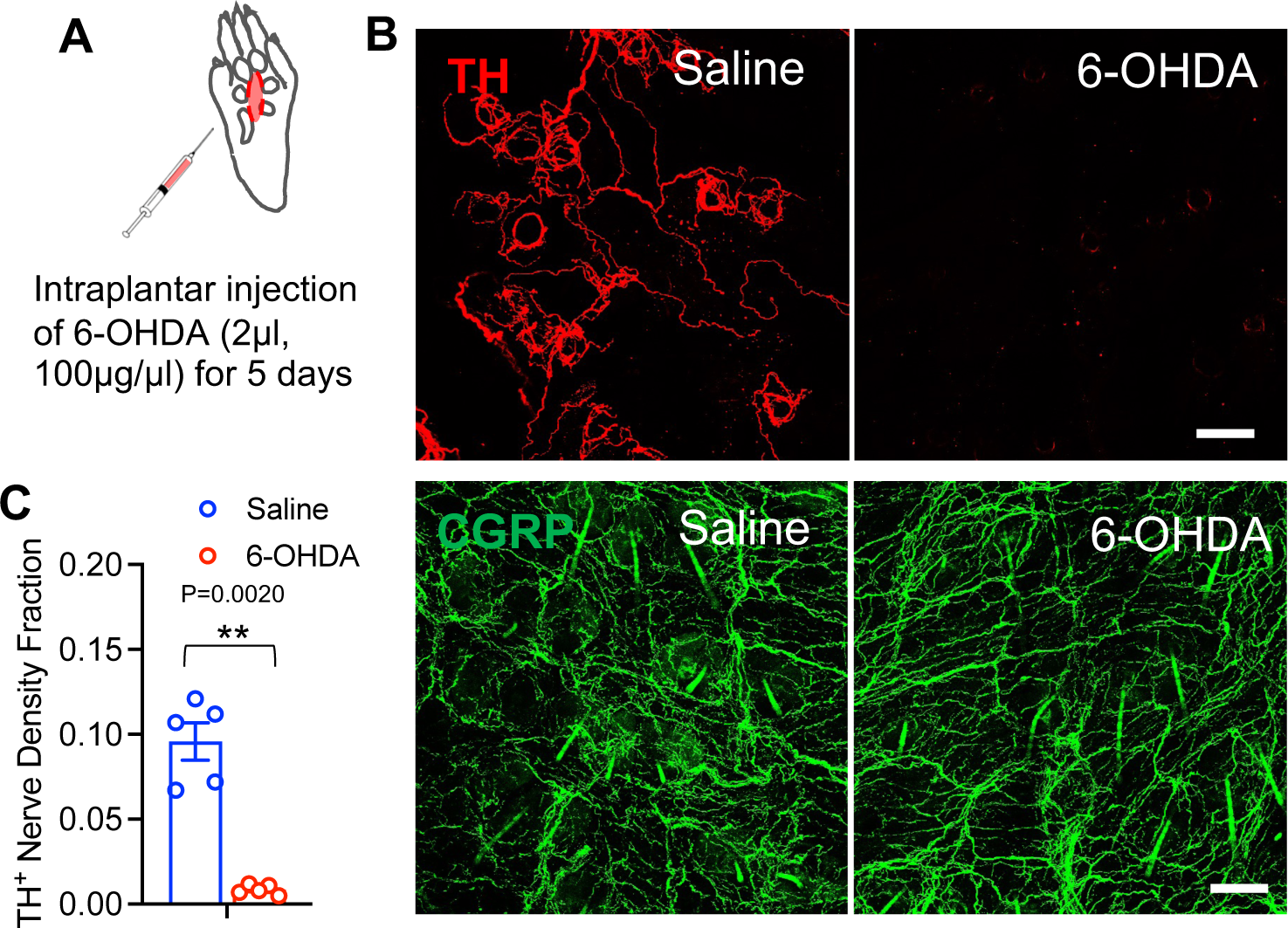
Local intraplantar injection of 6-OHDA eliminated TH^+^ nerve fibers in naïve mice. **(A)** Schematic for intraplantar injections of 6-OHDA (2μl, 100μg/μl in saline) for 5 consecutive days. **(B)** Whole mount immunostaining for TH (red) and CGRP (green) in the hind paw plantar hairy skin after local intraplantar injection of saline (left panel) or 6-OHDA (right panel) into C57BL6 mice. **(C)** Quantification of TH nerve density fraction in experiment shown in panel (B) (saline, blue, n=5; 6-OHDA, red, n=5). Data are presented as mean ± SEM. Paired two-tailed Student’s t-test. **P=0.0020, saline vs. 6-OHDA.

## References

Abdus-Saboor I, Fried NT, Lay M, Burdge J, Swanson K, Fischer R, Jones J, Dong P, Cai W, Guo X, Tao YX, Bethea J, Ma M, Dong X, Ding L, Luo W (2019) Development of a mouse pain scale using sub-second behavioral mapping and statistical modeling. Cell Rep 28:1623–1634.e4.

Abraira VE et al (2017) The cellular and synaptic architecture of the mechanosensory dorsal horn. Cell 168:295–310.e19.

Ahcan U, Arnez ZM, Bajrovic F, Janko M (1998) Contribution of collateral sprouting to the sensory and sudomotor recovery in the human palm after peripheral nerve injury. Br J Plast Surg 51:436–443.

Ali Z, Meyer RA, Belzberg AJ (2002) Neuropathic pain after C7 spinal nerve transection in man. Pain 96:41–47.

Arcourt A, Gorham L, Dhandapani R, Prato V, Taberner FJ, Wende H, Gangadharan V, Birchmeier C, Heppenstall PA, Lechner SG (2017) Touch receptor-derived sensory information alleviates acute pain signaling and fine-tunes nociceptive reflex coordination. Neuron 93:179–193.

Bai L, Lehnert BP, Liu J, Neubarth NL, Dickendesher TL, Nwe PH, Cassidy C, Woodbury CJ, Ginty DD (2015) Genetic identification of an expansive mechanoreceptor sensitive to skin stroking. Cell 163:1783–1795.

Basbaum AI, Bautista DM, Scherrer G, Julius D (2009) Cellular and molecular mechanisms of pain. Cell 139:267–284.

Bourquin AF, Suveges M, Pertin M, Gilliard N, Sardy S, Davison AC, Spahn DR, Decosterd I (2006) Assessment and analysis of mechanical allodynia-like behavior induced by spared nerve injury (SNI) in the mouse. Pain 122:14.e1–14.14.

Campbell JN, Meyer RA (2006) Mechanisms of neuropathic pain. Neuron 52:77–92.

Cobianchi S, de Cruz J, Navarro X (2014) Assessment of sensory thresholds and nociceptive fiber growth after sciatic nerve injury reveals the differential contribution of collateral reinnervation and nerve regeneration to neuropathic pain. Exp Neurol 255:1–11.

Collyer E, Catenaccio A, Lemaitre D, Diaz P, Valenzuela V, Bronfman F, Court FA (2014) Sprouting of axonal collaterals after spinal cord injury is prevented by delayed axonal degeneration. Exp Neurol 261:451–461.

Costigan M, Scholz J, Woolf CJ (2009) Neuropathic pain: A maladaptive response of the nervous system to damage. Annu Rev Neurosci 32:1–32.

Diamond J, Foerster A (1992) Recovery of sensory function in skin deprived of its innervation by lesion of the peripheral nerve. Exp Neurol 115:100–103.

Diamond J, Coughlin M, Macintyre L, Holmes M, Visheau B (1987) Evidence that endogenous beta nerve growth factor is responsible for the collateral sprouting, but not the regeneration, of nociceptive axons in adult rats. Proc Natl Acad Sci U S A 84:6596–6600.

Doucette R, Diamond J (1987) Normal and precocious sprouting of heat nociceptors in the skin of adult rats. J Comp Neurol 261:592–603.

Duraku LS, Hossaini M, Schuttenhelm BN, Holstege JC, Baas M, Ruigrok TJ, Walbeehm ET (2013) Re-innervation patterns by peptidergic substance-P, non-peptidergic P2X3, and myelinated NF-200 nerve fibers in epidermis and dermis of rats with neuropathic pain. Exp Neurol 241:13–24.

Duraku LS, Hossaini M, Hoendervangers S, Falke LL, Kambiz S, Mudera VC, Holstege JC, Walbeehm ET, Ruigrok TJ (2012) Spatiotemporal dynamics of re-innervation and hyperinnervation patterns by uninjured CGRP fibers in the rat foot sole epidermis after nerve injury. Mol Pain 8:61–61.

Gangadharan V, Zheng H, Taberner FJ, Landry J, Nees TA, Pistolic J, Agarwal N, Mannich D, Benes V, Helmstaedter M, Ommer B, Lechner SG, Kuner T, Kuner R (2022) Neuropathic pain caused by miswiring and abnormal end organ targeting. Nature 606:137–145.

Ghitani N, Barik A, Szczot M, Thompson JH, Li C, Le Pichon CE, Krashes MJ, Chesler AT (2017) Specialized mechanosensory nociceptors mediating rapid responses to hair pull. Neuron 95:944–954.e4.

Gloster A, Diamond J (1995) NGF-dependent and NGF-independent recovery of sympathetic function after chemical sympathectomy with 6-hydroxydopamine. J Comp Neurol 359:586–594.

Harrison BJ, Venkat G, Hutson T, Rau KK, Bunge MB, Mendell LM, Gage FH, Johnson RD, Hill C, Rouchka EC, Moon L, Petruska JC (2015) Transcriptional changes in sensory ganglia associated with primary afferent axon collateral sprouting in spared dermatome model. Genom Data 6:249–252.

Horch K (1981) Absence of functional collateral sprouting of mechanoreceptor axons into denervated areas of mammalian skin. Exp Neurol 74:313–317.

Jackson PC, Diamond J (1984a) Temporal and spatial constraints on the collateral sprouting of low-threshold mechanosensory nerves in the skin of rats. J Comp Neurol 226:336–345.

Jackson PC, Diamond J (1984b) Temporal and spatial constraints on the collateral sprouting of low-threshold mechanosensory nerves in the skin of rats. J Comp Neurol 226:336–345.

Jeon SM, Chang D, Geske A, Ginty DD, Caterina MJ (2021) Sex-dependent reduction in mechanical allodynia in the sural-sparing nerve injury model in mice lacking merkel cells. J Neurosci 41:5595–5619.

Ji RR, Strichartz G (2004) Cell signaling and the genesis of neuropathic pain. Sci STKE 2004:reE14.

Kim YS, Chu Y, Han L, Li M, Li Z, LaVinka PC, Sun S, Tang Z, Park K, Caterina MJ, Ren K, Dubner R, Wei F, Dong X (2014) Central terminal sensitization of TRPV1 by descending serotonergic facilitation modulates chronic pain. Neuron 81:873–887.

Kostrzewa RM, Jacobowitz DM (1974) Pharmacological actions of 6-hydroxydopamine. Pharmacol Rev 26:199–288.

Lakatos S, Jancso G, Horvath A, Dobos I, Santha P (2020) Longitudinal study of functional reinnervation of the denervated skin by collateral sprouting of peptidergic nociceptive nerves utilizing laser doppler imaging. Front Physiol 11:439.

Latremoliere A et al (2018) Neuronal-specific TUBB3 is not required for normal neuronal function but is essential for timely axon regeneration. Cell Rep 24:1865–1879.e9.

Lemaitre D, Hurtado ML, De Gregorio C, Onate M, Martinez G, Catenaccio A, Wishart TM, Court FA (2020) Collateral sprouting of peripheral sensory neurons exhibits a unique transcriptomic profile. Mol Neurobiol 57:4232–4249.

Li L, Rutlin M, Abraira VE, Cassidy C, Kus L, Gong S, Jankowski MP, Luo W, Heintz N, Koerber HR, Woodbury CJ, Ginty DD (2011) The functional organization of cutaneous low-threshold mechanosensory neurons. Cell 147:1615–1627.

Livingston WK (1947) Evidence of active invasion of denervated areas by sensory fibers from neighboring nerves in man. J Neurosurg 4:140–145.

Mackinnon SE, Colbert SH (2008) Nerve transfers in the hand and upper extremity surgery. Tech Hand Up Extrem Surg 12:20–33.

Michel PP, Hefti F (1990) Toxicity of 6-hydroxydopamine and dopamine for dopaminergic neurons in culture. J Neurosci Res 26:428–435.

Morelli C, Castaldi L, Brown SJ, Streich LL, Websdale A, Taberner FJ, Cerreti B, Barenghi A, Blum KM, Sawitzke J, Frank T, Steffens LK, Doleschall B, Serrao J, Ferrarini D, Lechner SG, Prevedel R, Heppenstall PA (2021a) Identification of a population of peripheral sensory neurons that regulates blood pressure. Cell Rep 35:109191.

Morelli C, Castaldi L, Brown SJ, Streich LL, Websdale A, Taberner FJ, Cerreti B, Barenghi A, Blum KM, Sawitzke J, Frank T, Steffens LK, Doleschall B, Serrao J, Ferrarini D, Lechner SG, Prevedel R, Heppenstall PA (2021b) Identification of a population of peripheral sensory neurons that regulates blood pressure. Cell Rep 35:109191.

Nascimento FP, Magnussen C, Yousefpour N, Ribeiro-da-Silva A (2015) Sympathetic fibre sprouting in the skin contributes to pain-related behaviour in spared nerve injury and cuff models of neuropathic pain. Mol Pain 11:59-x.

Nath RK, Mackinnon SE (2000) Nerve transfers in the upper extremity. Hand Clin 16:131–9, ix.

Qu L, Caterina MJ (2016) Enhanced excitability and suppression of A-type K(+) currents in joint sensory neurons in a murine model of antigen-induced arthritis. Sci Rep 6:28899.

Rigoni M, Negro S (2020) Signals orchestrating peripheral nerve repair. Cells 9:10.3390/cells9081768.

Rutlin M, Ho CY, Abraira VE, Cassidy C, Bai L, Woodbury CJ, Ginty DD (2014) The cellular and molecular basis of direction selectivity of adelta-LTMRs. Cell 159:1640–1651.

Theriault M, Dort J, Sutherland G, Zochodne DW (1998) A prospective quantitative study of sensory deficits after whole sural nerve biopsies in diabetic and nondiabetic patients. surgical approach and the role of collateral sprouting. Neurology 50:480–484.

Usoskin D, Furlan A, Islam S, Abdo H, Lonnerberg P, Lou D, Hjerling-Leffler J, Haeggstrom J, Kharchenko O, Kharchenko PV, Linnarsson S, Ernfors P (2015) Unbiased classification of sensory neuron types by large-scale single-cell RNA sequencing. Nat Neurosci 18:145–153.

Walcher J, Ojeda-Alonso J, Haseleu J, Oosthuizen MK, Rowe AH, Bennett NC, Lewin GR (2018) Specialized mechanoreceptor systems in rodent glabrous skin. J Physiol 596:4995–5016.

Woolf CJ, Ma Q (2007) Nociceptors--noxious stimulus detectors. Neuron 55:353–364.

Xie W, Strong JA, Zhang JM (2017) Active nerve regeneration with failed target reinnervation drives persistent neuropathic pain. eNeuro 4:10.1523/ENEURO.0008-Feb.

Zheng Q, Xie W, Luckemeyer DD, Lay M, Wang XW, Dong X, Limjunyawong N, Ye Y, Zhou FQ, Strong JA, Zhang JM, Dong X (2022) Synchronized cluster firing, a distinct form of sensory neuron activation, drives spontaneous pain. Neuron 110:209–220.e6.

